# Multiple quantitative trait loci contribute tolerance to bacterial canker incited by *Pseudomonas syringae* pv. *actinidiae* in kiwifruit (*Actinidia chinensis*)

**DOI:** 10.1101/526798

**Authors:** Jibran Tahir, Stephen Hoyte, Heather Bassett, Cyril Brendolise, Abhishek Chatterjee, Kerry Templeton, Cecilia Deng, Ross Crowhurst, Mirco Montefiori, Ed Morgan, Andrew Wotton, Keith Funnell, Claudia Wiedow, Mareike Knaebel, Duncan Hedderley, Joel Vanneste, John McCallum, Kirsten Hoeata, David Chagné, Luis Gea, Susan E. Gardiner

**Affiliations:** The New Zealand Institute for Plant & Food Research Limited, Private Bag 11030, Manawatu Mail Centre, Palmerston North, 4442, New Zealand; The New Zealand Institute for Plant & Food Research Limited, Hamilton, New Zealand; The New Zealand Institute for Plant & Food Research Limited, Private Bag 92–169, Auckland, 1025, New Zealand; New Plant Soc. Cons. Agr., via Malpighi 5, Forlì, 47122, Italy; The New Zealand Institute for Plant & Food Research Limited, Lincoln, New Zealand; The New Zealand Institute for Plant & Food Research Limited, 412 No 1 Road, RD2, Te Puke 3182, New Zealand

**Keywords:** Psa, kiwifruit, QTLs, field tolerance, bioassays, oligogenic tolerance, innate immunity

## Abstract

*Pseudomonas syringae* pv. *actinidiae* (Psa) Biovar 3, a virulent, canker-inducing pathogen is an economic threat to the kiwifruit (*Actinidia* spp.) industry worldwide. The commercially grown diploid (2*x*) *A. chinensis* var. *chinensis* is more susceptible to Psa than tetraploid and hexaploid kiwifruit. However information on the genetic loci modulating *Psa* resistance in kiwifruit is not available. Here we report mapping of quantitative trait loci (QTLs) regulating tolerance to Psa in a diploid kiwifruit population, derived from a cross between an elite Psa-susceptible ‘Hort16A’ and a tolerant male breeding parent P1. Using high-density genetic maps and intensive phenotyping, we identified a single QTL for Psa tolerance on Linkage Group (LG) 27 of ‘Hort16A’ revealing 16-19% phenotypic variance and candidate alleles for susceptibility and tolerance at this loci. In addition, six minor QTLs were identified in P1 on distinct LGs, exerting 4-9% variance. Complete tolerance in the F1 population is attained by additive effects from ‘Hort16A’ and P1 QTLs providing evidence that divergent genetic pathways fend-off virulent Psa strain. Two different bioassays further identified new QTLs for tissue-specific responses to Psa. Transcriptome analysis of Psa-tolerant and susceptible genotypes in field revealed hallmarks of basal defense and provided candidate RNA-biomarkers for screening Psa tolerance.

## Introduction

*Pseudomonas syringae* is a hemi-biotrophic bacterial complex ^1^ that can infect a range of plant species. It comprises pathovars which cause similar symptoms on their host plants and several pathovars can lead to severe crop loss. *P. syringae* pv. *actinidiae* (Psa) infects several species of *Actinidia* (kiwifruit) ^2,3^ and virulent Psa strains induce a range of symptoms on the main stem of the vine, foliage, floral buds and fruits ^4^. Psa pathovar’s strains can be grouped into five biovars based on their genetic and biological characteristics ^4,5^. Strains of *biovar* 3, previously called Psa*-*V (referred to here as Psa), are currently the most aggressive and were responsible for outbreaks from the year 2008 ^6–9^. Psa has cost the kiwifruit industry billions of dollars worldwide and its incursion in New Zealand in 2010 completely destroyed vines of the Psa-susceptible diploid *A. chinensi*s ‘Hort16A’ ^4,8,10^.

Most of the globally cultivated cultivars of kiwifruit, including *A. chinensis* (A Planch.) var. *chinensis*, *A. chinensis* (A Chev.) C.F. Liang *et* A.R. Ferguson var. *deliciosa*, as well as accessions from *A. arguta* and *A. kolomikta* are natural hosts of Psa ^10–18^. Early reports of Psa infections and symptoms in *Actinidia* species emerged from Japan, China, Korea and Italy from 1984 to 1994 ^2,3,13,14,16,17,19^. The symptoms include cankers on trunk and leaders, cane death and stem collapse, discharge of red and milky exudates (ooze) from cankers, canes and abaxial leaf surfaces, tip browning, angular leaf necrosis (sometimes with chlorotic halos), shoot and leaf wilt, bud browning and flower blight. Strains of Psa infect *Actinidia* species with varying degrees of virulence, indicating a classical host-pathogen evolutionary relationship ^5,9,20–24^.

Screening of thousands of *Actinidia* genotypes from 24 taxa in the breeding program at The New Zealand Institute for Plant & Food Research Limited (PFR) for tolerance to natural and artificial Psa infections ^25,26^ revealed that diploid (2*x*) *A. chinensis* var. *chinensis* are more susceptible to Psa infection than tetraploid (4*x*) *A. chinensis* var. *chinensis*, which in turn are more susceptible than diploid and hexaploid (6*x*) *A. chinensis* var. *deliciosa* ^25–27^. Many species outside the *A. chinensis* complex are more tolerant to Psa than *A. chinensis* and the germplasm holds diverse genetic potential for Psa tolerance ^28^. Information on the genetic markers and molecular mechanisms associated with *Psa* tolerance and resistance in the commercial cultivars producing taxas including *A. chinensis*, *A. deliciosa* and *A. arguta* is however limited. In this study we provide the first detailed view of the genetic loci modulating Psa tolerance and tissue-specific response in diploid *A. chinensis*, utilizing an intensively phenotyped population of seedlings developed from a cross between Psa-susceptible ‘Hort16A’ and a tolerant breeding parent (P1), as our experimental material for quantitative trait locus (QTL) analysis.

## Results

### Intensive phenotyping targets diverse developmental stages and environmental conditions

Initially, a pilot population of 53 genotypes from ‘Hort16A’ × P1 were replicated 3 times and phenotyped following natural field infection with Psa. The response to Psa infection in the expanded population was measured on 236 genotypes of ‘Hort16A’×P1 population, which were clonally replicated ~30 times. Phenotyping of the population was performed under field conditions following natural infection as well as using two bioassays (scheme for phenotyping is laid out in Supplementary Fig.1a). Multiple phenotypes were recorded in field (Fig. 1a-e) to develop a combined score referred to as Psa_score_Field (Fig. 2a). The mean clonal repeatability for this score was 0.65, while the repeatability of clonal means at 0.8. For the stab assay ^26^, various tissue-specific phenotypic responses were recorded, including Stem_necrosis, Leaf_spots, Ooze, Stem_collapse, Tip_death and Wilt (Fig. 1f-k, Fig. 2b), with repeatability of clonal means for these scores as 0.60, 0.766, 0.64, 0.79, 0.78 and 0.71 respectively. A Psa_score_Stab was also calculated (Fig. 2b) from all phenotypes assessed in the Stab assay (see Experimental Procedures). In the flood bioassay, adapted from ^29^, overall health was scored at weekly intervals post-inoculation (Flood Assay/FA_Week1 to FA_Week5) (Fig. 1l). The frequency distribution of phenotypes and Psa_scores revealed that most exhibited non-Normal distribution based on the Shapiro-Wilk test (Fig. 2). However, for the stab assay, the majority of the observations displayed normal distributions. (Fig. 2b). As such, the correlation among the phenotypic scores from the field assessments and the bioassays was found to be poor (Supplementary Fig.1b). The correlation among different phenotypes within the bioassays was medium to high. A 3-dimensional principal components analysis (PCA) on the correlation matrix of the field assessment, stab assay and flood assay displays a high degree of divergence in the rankings of the population for Psa response and tolerance when assessed through different approaches (Fig. 2d).

**Fig. 1.**
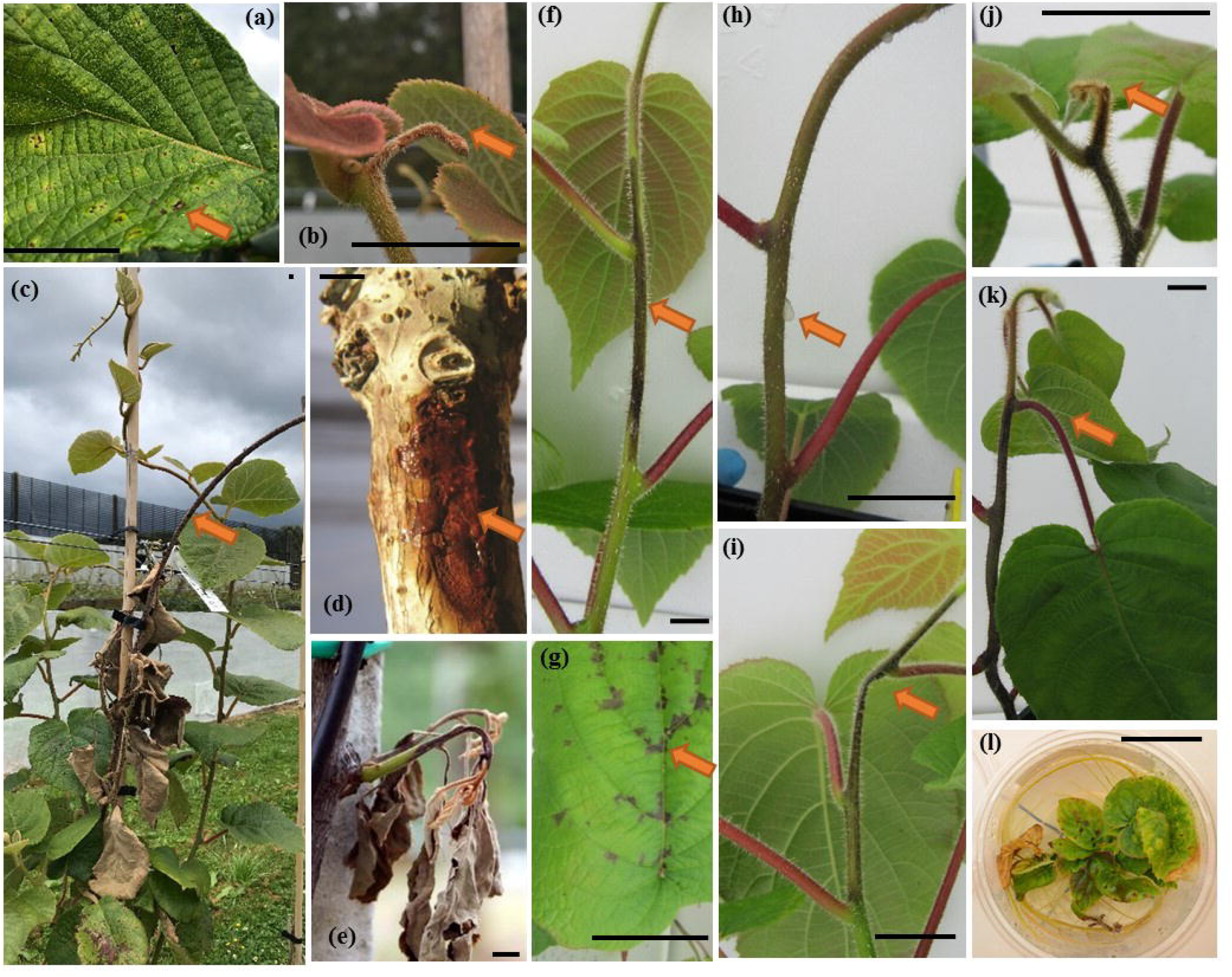
Phenotypic responses in *Actinidia chinensis* plants in response to *Pseudomonas syringae* pv. *actinidiae* (Psa) exposure. Field phenotypes include (a) Leaf_spots (b) Tip_death (c) Cane_death (d) Ooze and (e) Shoot_death. Phenotypes observed in the stab bioassay include f) Stem_necrosis, g) Leaf_spots, h) Ooze, i) Stem_collapse, j) Tip_death, k) Wilt and l) is a representative flood assay (FA_Week3) phenotype for disease response.

**Fig. 2.**
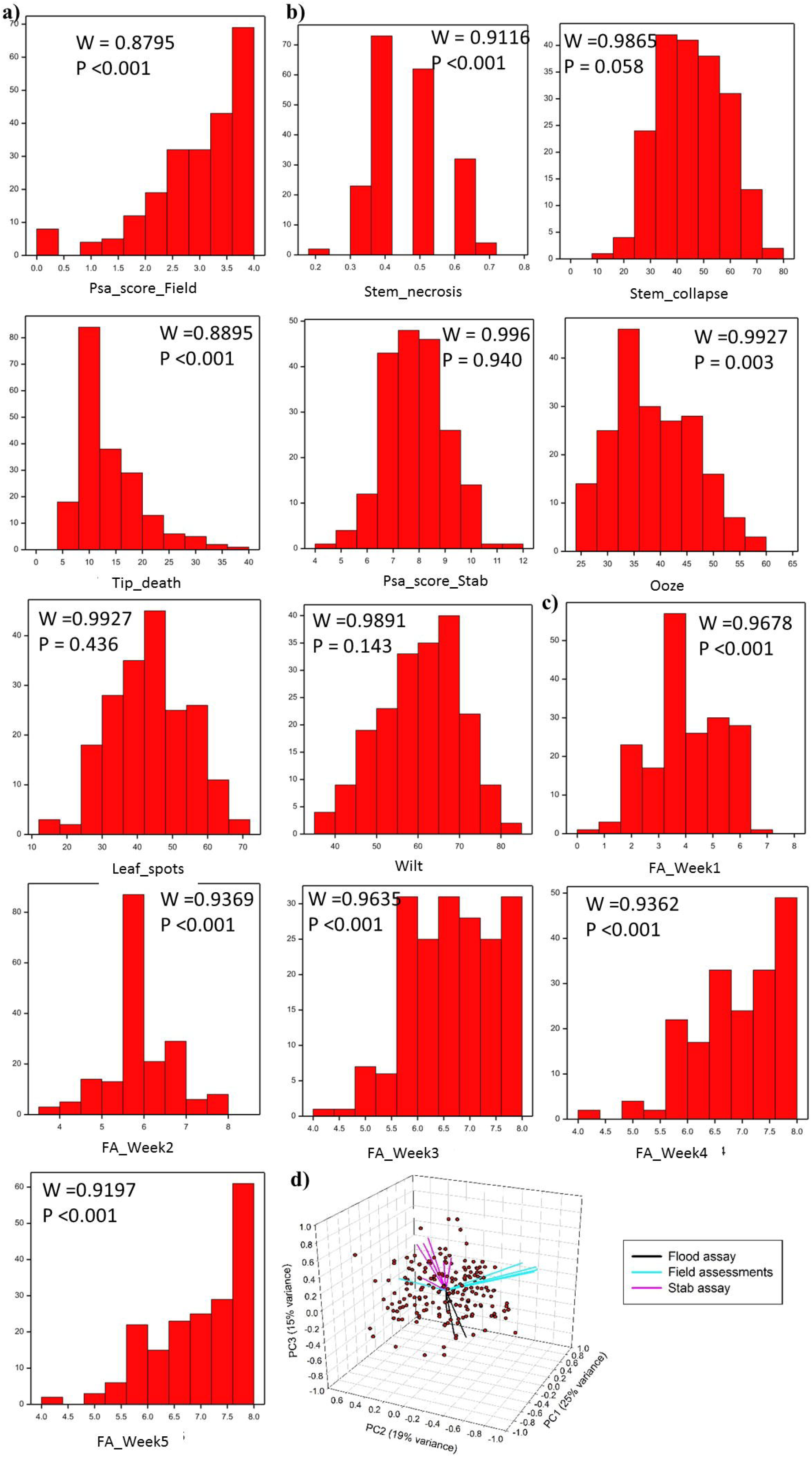
Distribution of phenotypes in the ‘Hort16A’ × P1 mapping population of 236 genotypes. a) Least squares mean (LS Mean) of Psa_score_Field after 15 months in the field. The x-axis displays the progression of susceptibility from left to right, while the y-axis represents frequency in the population. b) LSM of phenotypes from the stab assay, including Stem_necrosis, Stem_collapse, Tip_death, Psa_score_Stab, Ooze, Leaf_spot and Wilt. c) Means of the health score from Flood bioassays (FA_Week 1 to FA_Week5). The WSTATISTIC is from the Shapiro-Wilks test for the null hypothesis that the distribution is normal. Phenotypic scores with P<0.001 are rejected for the hypothesis that these distributions are normal. d) Principal components analysis on the correlation matrix of the field assessment, flood assay and stab assay measures. Genotypes are shown as points and measurements are shown as vectors (lines pointing from the origin) defined by their correlation with the three principal components.

### Genotyping-by-Sequencing provided high-density genetic maps for ‘Hort16A’×P1 genotypes

Using genotyping-by-sequencing (GBS)^30^, the population of ‘Hort16A’×P1 enabled the construction of high-density genetic maps utilizing 3,777 and 3,454 SNP markers, for ‘Hort16A’ and P1 respectively (Supplementary Fig. 2 and 3) using Red5^31^ and Hongyang^32^ as reference genomes. The maps for ‘Hort16A’ and P1 encompassed a total genetic distance of 3,499 cM and 3,875 cM, respectively, with an average density of 1 marker/ 2cM for both parents. All predicted 29 LGs were constructed for ‘Hort16A’; however some were fragmented in P1 (LGs 3, 16, 19, 23, 25, 27).

**Fig. 3.**
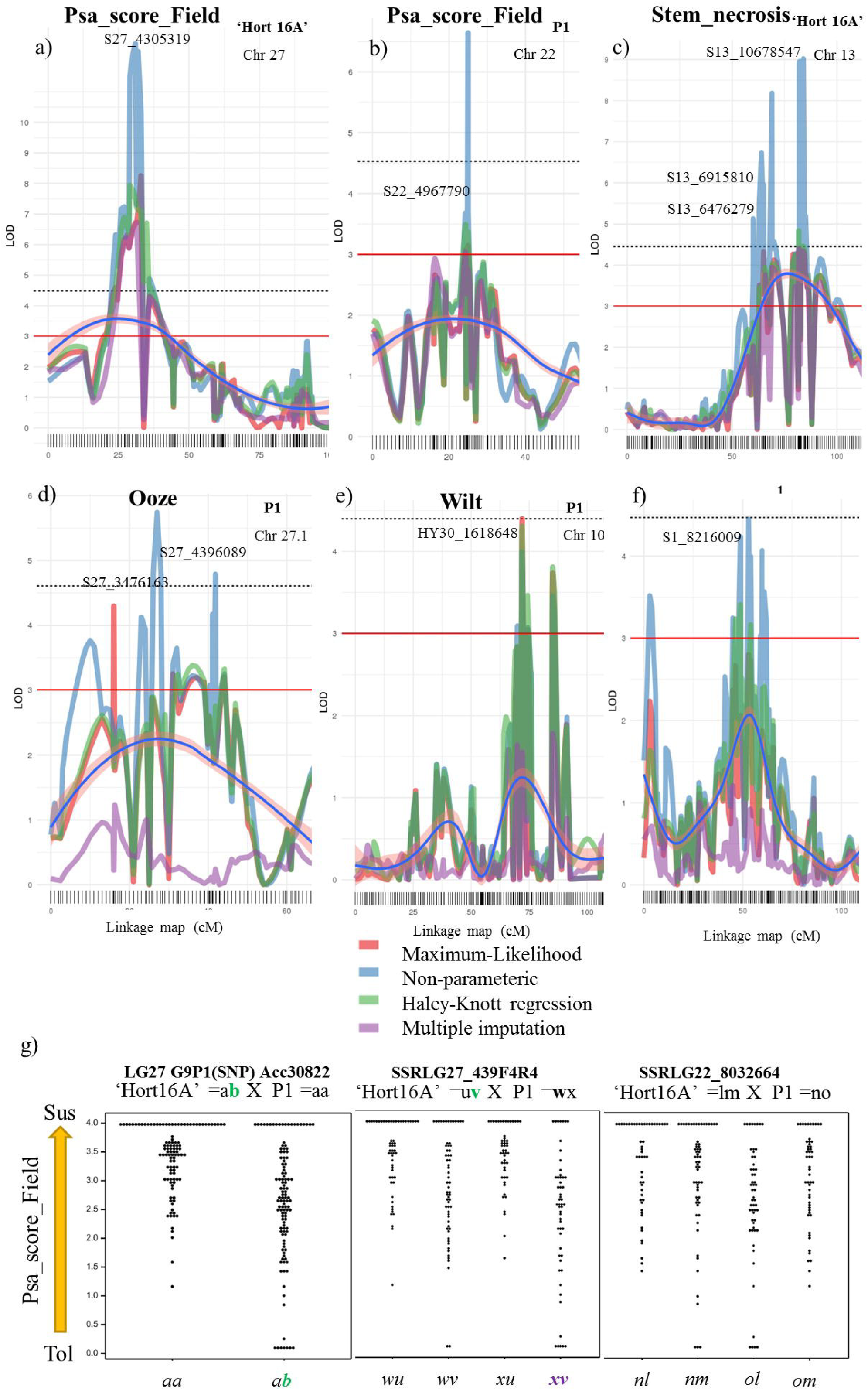
Quantitative trait loci (QTLs) from ‘Hort16A’ and P1 for control of field tolerance and tissue specific symptomatic responses to Psa. The outputs depict quantitative trait loci (QTL) scans with different models. a) linkage group (LG)27 of ‘Hort16A’, b) LG22 of P1, both for Psa_score_Field. From stab assay phenotypes major QTLs on: c) LG13 in ‘Hort16A’ for Stem_necrosis, and in P1 on d) the upper arm of LG27 for Ooze, e) LG10 for Wilt and f) LG1 for Psa_score_Stab. SNPs at peaks are indicated. g) shows dot plot analysis of the allelotypes of markers underlying quantitative trait loci derived from the Psa_score_Field for the population ‘Hort 16A’ x P1’.

### QTL mapping from field phenotype scores confirmed oligogenic nature of Psa field tolerance

A QTL for control of field tolerance to Psa, Psa_score_Field, was identified in ‘Hort16A’ on the upper arm of LG27 (Fig. 3a) using multiple models for QTL discovery. At a LOD score of 7.02 (Fig. 3a), the location of the LG27 QTL on the Red5 genome (version 1.69.0) ^33^ is between ~3.4 to 4.6 Mbp. The LG27 QTL was also identified for Psa_score-Field, in ‘Hort16A’ from the pilot trial (Supplementary Table 1). .A SNP marker G9P1 developed from Acc30822, a gene of unknown function underlying the QTL and a multi-allelic Simple Sequence Repeat (SSR) marker SSRLG27_439F4R4, contributed 16% (favorable allele *b*) and 19% (favorable allele *v*, band size 428 bp) of the population phenotypic variance, respectively (Fig. 3g and Supplementary Table 1). The multi-allelic SSR marker revealed the contribution of the favorable 428 bp *A. chinensis* grandparental allele *v*, to Psa tolerance (Fig. 3g), compared to the other 408 bp allele *u* which is associated with susceptibility.

**Table 1.**
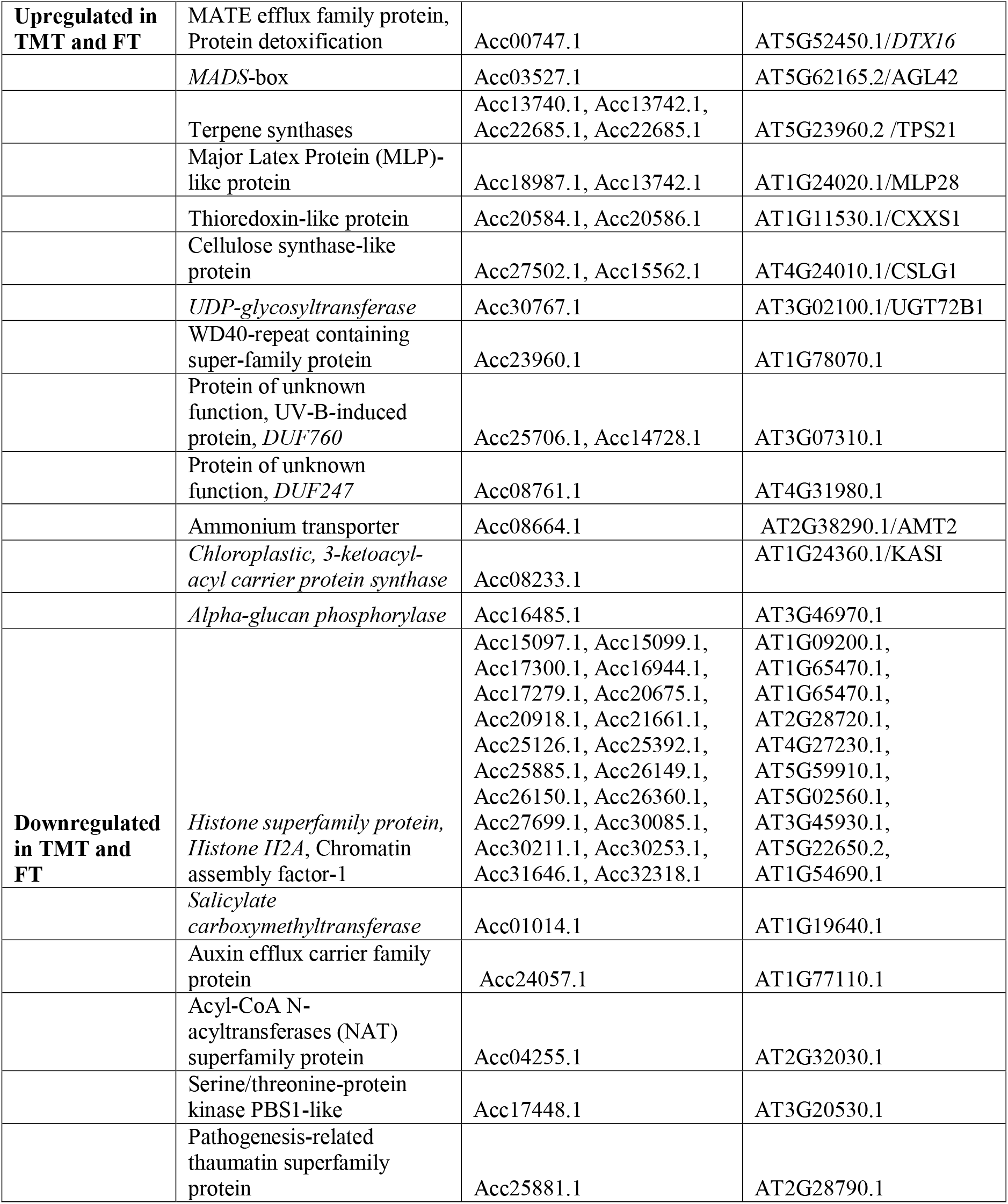
Candidates from differentially expressed genes in field tolerant genotypes. Psa-tolerant plants (FT), Psa Tolerant to Medium Tolerant /Psa-TMT and Psa-susceptible genotypes (Psa-Sus).

Using interval mapping and KW analysis, six QTLs were identified in P1, for Psa_score_Field indicating Psa tolerance is multigenic in P1. A single QTL with LOD score above 3 was located on the upper arm of LG22 (Fig. 3b), while three additional QTLs on LGs 3.1, 15 and 24 (Supplementary Fig. 4), as well as two KW QTLs on LG14 (S14_5310060, *K* value > 9, *P* < 0.0001) and LG28 (S28_1476180, *K* value > 7, *P* < 0.0001). From these, the effect of favorable grandparent alleles from P1 on field tolerance was verified from at least 3 QTLs i.e., by analysis of an SSR marker designed in the region underlying the LG22 QTL (SSRLG22_8032664) (Fig 3g and Supplementary Table 1), a SNP marker E6P3 designed in a putative cell wall protein encoding gene Acc15766 within the LG14 QTL (Supplementary Fig. 5, Supplementary Table 1), and the LG28 QTL (SSRLG28_1378F5R5) (Supplementary Fig. 5). Screening of SSRLG27_439F4R4 in another set of field-grown ‘Hort16A’ × P1 progeny confirmed association of the *v* allele with tolerance (Supplementary Fig. 6a). The combination of favorable alleles from the ‘Hort16A’ LG27 QTL and three QTLs from P1 (LGs 14, 22 and 28) yielded a percentage variance of 40.6% (Supplementary Table 1): this combination identified ~80% of the tolerant ‘Hort16A’ × P1 genotypes in the field (Supplementary Fig. 7).

**Fig. 4.**
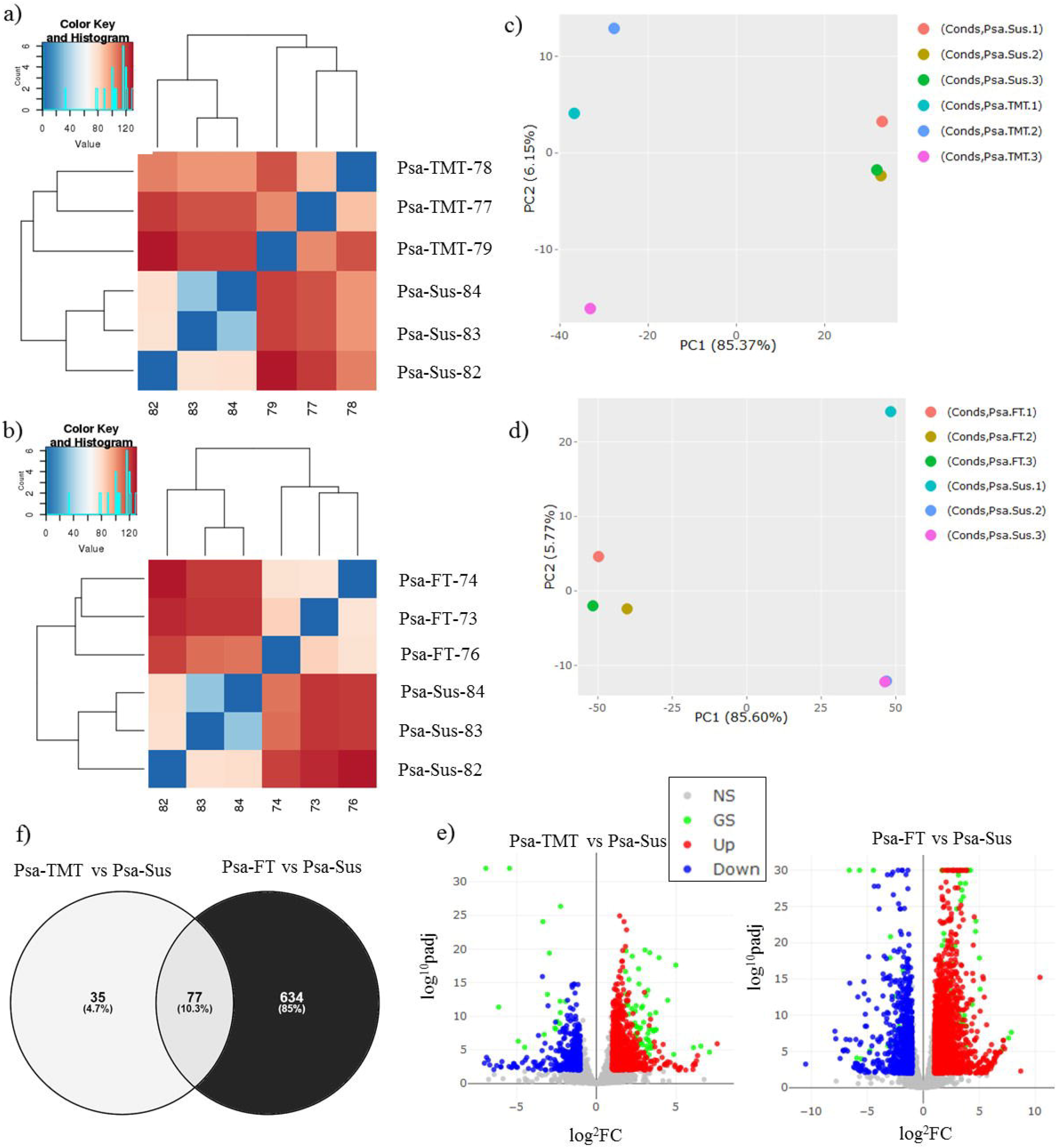
RNA-seq analysis of ‘Hort16A’ × P1 genotypes, exhibiting tolerance or susceptibility to Psa in the field. RNA-seq was performed on young healthy leaf tissues of field-grown plants belonging to three groups based on relative tolerance / susceptibility. The first group included three relatively Psa-tolerant plants (Tolerant to Medium Tolerant /Psa-TMT). The second group included three fully Psa-susceptible genotypes, including ‘Hort16A’ (Psa-Sus). All had been exposed to Psa for 1 year in the field. The third group represents the three most tolerant genotypes, tolerant over 3 years in the field (Psa-FT). a) and b) show heat-maps for the genome wide differential expression (DE) analysis in Psa-TMT vs Psa-Sus and Psa-FT vs Psa-Sus respectively. c and d) are plots of Principal component analysis for the DE in Psa-TMT vs Psa-Sus and Psa-FT vs Psa-Sus respectively. e) shows volcano plots for the DE, with significantly (Padj <0.01/log^10^ padj, logfold2) upregulated and downregulated genes highlighted in red and blue respectively in the two comparisons, Psa-TMT vs Psa-Sus and Psa-FT vs Psa-Sus. The grey dots indicate non-significantly expressed genes, whereas the green dots highlight the genes that are differentially expressed in common between Psa-TMT vs Psa-Sus and Psa-FT vs Psa-Sus. f) shows the DE genes in common or unique, respectively, between Psa-TMT vs Psa-Sus and Psa-FT vs Psa-Sus.

**Fig. 5.**
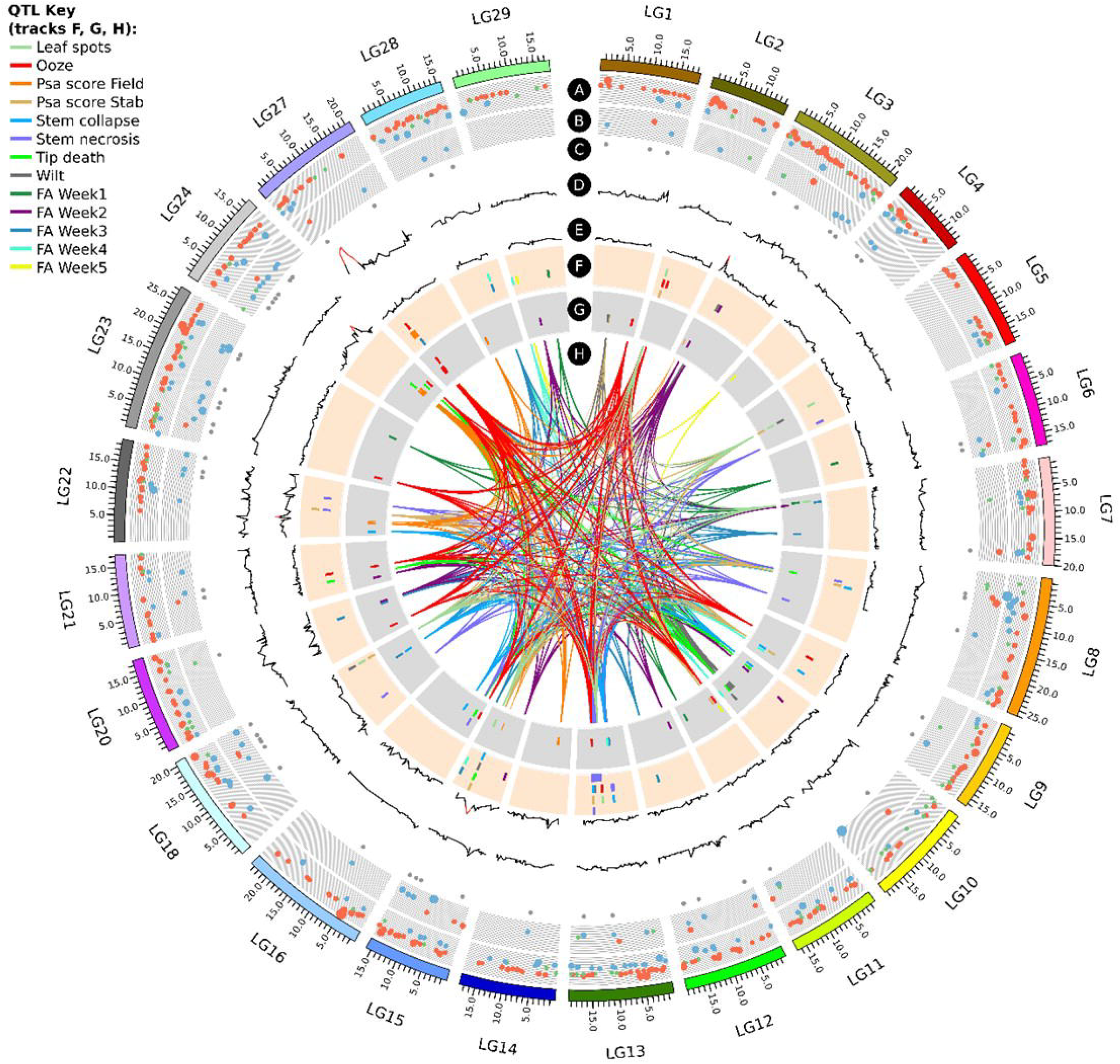
Circos plot of quantitative trait loci (QTLs) for various phenotypes in field and bioassay as well as RNA-seq data associated with Psa-tolerant and susceptible genotypes, anchored on the chromosomes of the Red5 genome version 1.69.0. Tracks A and B represent differentially expressed genes (DEGs) with logFC +2 and above in fully tolerant (Psa-FT) vs susceptible (Psa-Sus) and tolerant to medium tolerant (PsaTMT) vs Psa-Sus genotypes, respectively. On track A, blue circles are upregulated and red circles are downregulated genes in Psa-FT compared to Psa-Sus genotypes. On track B, blue circles are downregulated and red circles are upregulated genes in Psa-TMT compared to Psa-Sus genotypes. Genes with logFC, between 1 and −1, are represented by green circles. Increase in circle diameter indicates increasing logFC value. Track C represents DEGs common to Psa-FT and Psa-TMT. Track D and E are LOD values for Psa_score_Field for ‘Hort16A’ and P1 respectively. The lines change from black to red for a LOD score >3. Track F and G are QTLs detected from all phenotypes listed in QTL key in ‘Hort16A’ and P1 respectively. H represents the lines connecting QTLs for similar phenotype on different chromosomes.

**Fig. 6.**
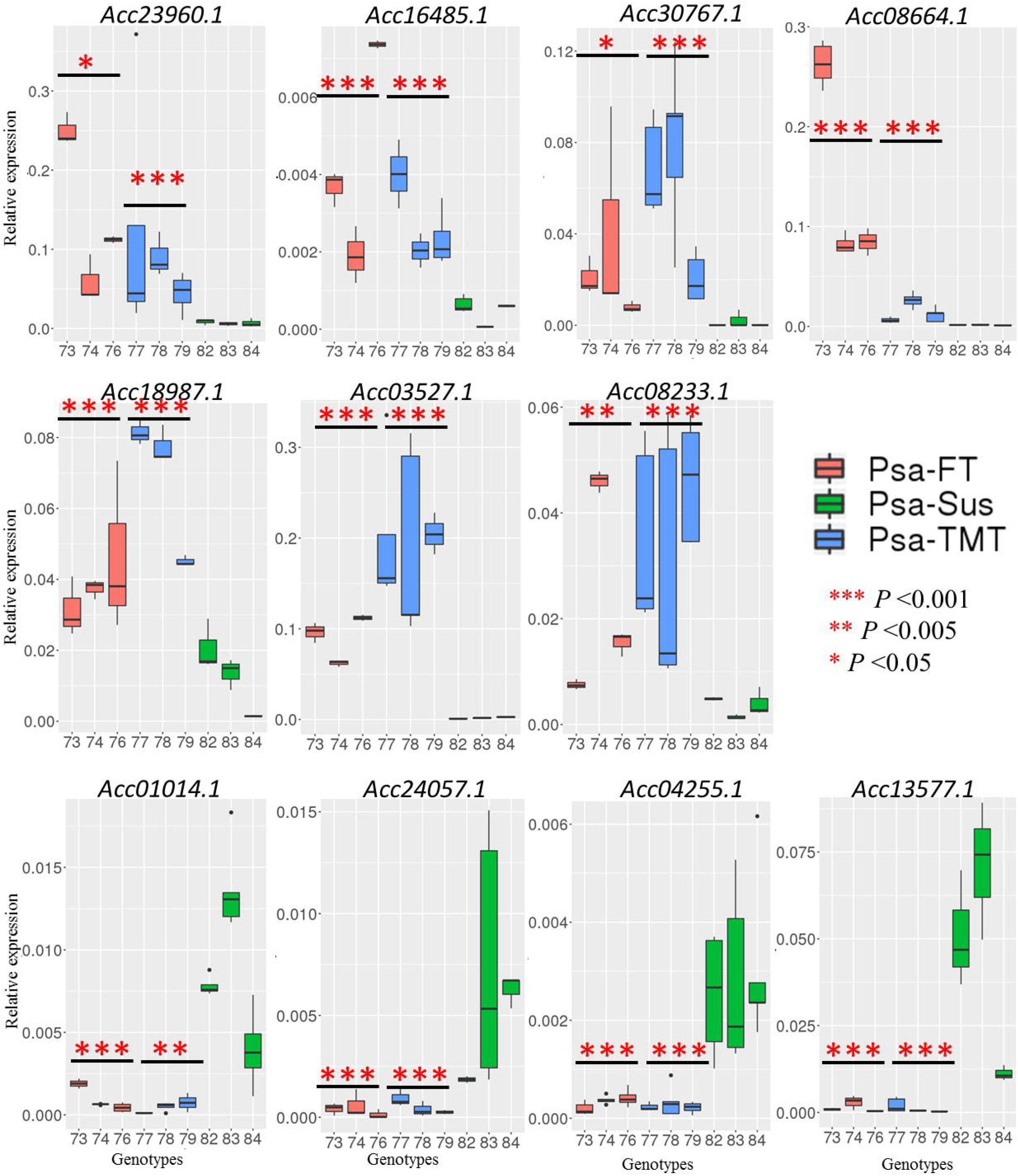
Real-time quantitative PCR of relative gene expression of candidate genes in Psa-TMT, Psa-FT and Psa-Sus genotypes. Expression of the candidate genes was analyzed in Psa-TMT (73, 74, 76), Psa-FT (77, 78, 79) and Psa-Sus (82, 83, 84) genotypes using real-time quantitative reverse transcription polymerase chain reaction (RT-qRT-PCR). Data represents mean relative gene expression of the candidate genes, in three to seven clonal replicates for each genotype, to the mean of *Actin* and *Ubiquitin* genes and plotted using geom_boxplot. Asterisks represents statistically significant differences in the relative expression of the candidate genes in genotypes of Psa-TMT and Psa-FT compared to the mean relative expression of genes in Psa-Sus genotypes using Student’s *t*-test.

Since the QTL identified in ‘Hort16A’ has the highest effect, we predicted that this locus could be linked to susceptibility observed in diploid kiwifruit breeding parents. For this purpose, we performed validation of the LG27 QTL in another field grown *A. chinensis* population, derived from two Psa-susceptible parents‘Hort22D’ and P2, using SSRLG27_439F5R5 located ~3 kb distant from SSRLG27_439F4R4. Data revealed that the 430/432 bp allele linked in *cis* with the previously described allele *v* was associated with tolerance in this population and was present in both parents, while the 440 bp allele *z* (size 440 bp) was linked to susceptibility and present in ‘Hort22D’ (Supplementary Fig. 6b). The region underlying the QTL on chromosome 27 spans orthologues of genes with putative functions involved in Pathogen-Associated Molecular Pattern (PAMPs)-triggered immunity (PTI), including cysteine rich receptor-like protein kinases, serine-threonine like protein kinase, enzymes involved in amino acid, carbohydrate and phosphate metabolism, sugar transporter, transcription factors, auxin-responsive and transport protein, epigenetic modulators and heat repeat-containing protein (Supplementary Table 2).

### Additional genetic hotspots associated with tissue and environment-specific phenotypic responses to Psa infection identified using bioassays

#### Analysis of Psa tolerance in P1 using stab assay and leaf infection

The stab assay targets the vascular system and enabled a range of different phenotypes to be scored following Psa infection (Fig. 1f to k). P1 appeared to be relatively tolerant in comparison with ‘Hort16A’ in the stab assay, as in the field and grouped close to Psa-tolerant *A. arguta* and *A. chinensis* var. *deliciosa* for the Stem_necrosis response to infection (Supplementary Fig. 8). Consistent with this, ‘Hort16A’ hosted significant growth of endophytic populations of Psa in the leaves, 10 days post-inoculation (Supplementary Fig. 9), compared with P1 and a Psa-tolerant tetraploid *A. chinensis* genotype both of which did not support endophytic growth of Psa over the same time period (Supplementary Fig. 9).

#### QTLs for control of stem necrosis and collapse, tip death and Psa score determined in the stab bioassay

Multiple interval mapping methods identified QTLs for control of Stem_necrosis on LG13 in ‘Hort16A’ at three positions; S13_6915810 and S13_10678547, with a LOD score ranging between 4.5 and 9 (Fig. 3c) and S13_13629983 (Supplementary Table 3). Moreover, QTLs were detected in the same region on LG13 for control of Stem_collapse and Psa_score_Stab, indicating these were genetic hotspots for host-pathogen interaction in vascular tissues. Interestingly, QTLs for the control of Stem_necrosis in P1 were identified on different chromosomes from those of ‘Hort16A’, namely the upper arm of LG16 and lower arm of LG23 (Supplementary Table 4 and Supplementary Fig. 10). As for ‘Hort16A’, QTLs from P1 coincided with those for other phenotypes including Tip_death and Psa_score_Stab. A significant QTL for control of Psa_score_Stab was also detected on LG1 of P1 (Fig. 3f, Supplementary Fig. 11). It was noticeable that the Tip_death phenotype generated multiple putative QTLs from both ‘Hort16A’ and P1 (Supplementary Fig. 12 and 13).

#### Oozing as a symptom of Psa infection

Oozing of a bacterial exudate was observed following Psa infection and QTLs for control of this phenotype were identified on LGs 2, 13 and LG15 (Supplementary Table 3 and Supplementary Fig.10) of ‘Hort16A’. For P1, QTLs were detected on the upper arm of LG27 (Fig 3d and Supplementary Table 4) and LG13. QTLs for control of the Ooze phenotype detected on LG13 and 27 overlapped QTLs detected in ‘Hort16A’ for the Stem_necrosis phenotype, as well as Psa_score_Field. Other QTLs identified in ‘Hort16A’ and P1 using KW analysis for the Ooze phenotype are listed in Supplementary Table 3 and 4, respectively.

#### Leaf spots and Wilt

We observed symptomatic responses to Psa infection in leaf tissues distant from the point of inoculation in the stem. In ‘Hort16A’, QTLs for Leaf_spots (Supplementary Table 3 and Supplementary Fig. 10) were detected on LGs 2, 5, 13 and 26. QTLs for Wilt in ‘Hort16A’were detected on LGs 3, 13, 15 and 18. Most of these overlapped QTLs identified for Ooze and Stem_necrosis. In P1, QTLs for Leaf_spots (Supplementary Table 4 and Supplementary Fig. 11) were detected on LGs 1 and 5. A significant QTL was detected on LG10 of P1 for Wilt (Fig. 3e and Supplementary Table 4).

#### Phenotypic tolerance to Psa exposure in tissue culture

When ‘Hort16A’ x P1 population grown aseptically in tissue culture were challenged with Psa, multiple QTLs were detected for a health score at each weekly time-point (FA_Week1 to FA_Week5) (Supplementary Table 5). For ‘Hort16A’, *K* values were significant on LG15 at the third and fourth weeks following infection. A QTL on LG27 with lower significance overlapped the major QTL on LG27 identified in ‘Hort16A’ for Psa_score_Field. For P1, a significant QTL identified on the upper arm of LG13 for 3 and 4 weeks post-infection and overlapped the QTL region identified from phenotypes in the stab assay. Plant phenotypes changed dramatically during the period post-infection and additional QTLs were identified for health score at different time points (Supplementary Table 5).

The coordinates for all the QTLs in the Red5 genome versions 1.69.0 ^33^ and 1.68.5 ^31^ are provided in Supplementary Data 1.

### RNA-seq of ‘Hort16A’, P1 and F1 genotypes exhibiting Psa tolerance or susceptibility in the field revealed patterns of innate immunity

RNA-seq performed on healthy young leaf tissues from three groups of ‘Hort16A’ × P1 F1 genotypes differing in field tolerance to Psa demonstrated clear differences in gene expression. The first group included three relatively tolerant-to medium-tolerant genotypes, including P1 (Psa-TMT), while the second group included three fully susceptible genotypes, including ‘Hort16A’ (Psa-Sus). At the same time, samples were harvested from the three most tolerant ‘Hort16A’xP1 genotypes, which had shown tolerance for four years in the field (Psa-FT). Heat maps and PCA plots of expression data from the pair-wise comparison between the three groups demonstrated extreme variation between the susceptible (Psa-Sus) and two tolerant groups (Psa-TMT and Psa-FT) (Fig. 4a to d). Differential gene expression analysis conducted between the groups of tolerant and susceptible genotypes at α<0.005 with *p* values adjusted < 0.1 revealed that from 31,588 genes, 23 (0.076%) were upregulated and 88 (0.28%) were downregulated in Psa-TMT compared with Psa-Sus (Supplementary Data2). Psa-FT genotypes exhibited 712 differentially expressed genes (DEGs) when compared with Psa-Sus. Of these, 172 (0.59%) were upregulated and 539 (1.9%) were downregulated in Psa-FT genotypes compared to Psa-Sus (Supplementary Data2). Seventy-seven genes (0.24%) were differentially expressed in common among tolerant genotypes of the Psa-FT and Psa-TMT groups when each was compared with Psa-Sus (Fig. 4e and f and Supplementary Data 2).

#### DEGs in Psa-TMT and Psa-FT groups compared to Psa-Sus genotypes

The gene families upregulated in common in Psa-TMT and Psa-FT genotypes are mostly orthologues of protein-coding genes involved in plant basal defense against pathogens or Pathogen-Associated Molecular Patterns (PAMPs)-triggered immunity (PTI), cost of defense, cell wall and carbohydrate metabolism and other functions (Table 1). Psa-FT genotypes exhibit upregulation of a high number of genes with functions related to defense. The genes significantly downregulated in common in both tolerant genotypes, Psa-TMT and Psa-FT compared with Psa-Sus, are orthologues of protein coding genes involved in chromatin modulation such as histone encoding proteins, auxin efflux, and abiotic and biotic defense (Table 1).

#### Integrated view of QTLs and DEG in field

A Circos plot of all the QTLs and the DEGs anchored on the Red5 genome 1.69.0, highlighted a number of DEGs that co-localized with the QTL regions (Fig. 5). Circos diagrams for individual phenotypes are presented in Supplementary Figs.14 to 16 for phenotypes from the field, Stab bioassay and Flood bioassay, respectively.

#### Validation of expression of candidate genes

From the list of candidate DEGs (Table 1), relative expression of a few genes with diverse putative functions was verified in the genotypes from all three groups (Psa-TMT, Psa-FT, Psa-Sus), using real time quantitative reverse transcription polymerase chain reaction (RT-qRT-PCR) (Fig 6). Genes including Acc23960.1 (*Transducin/WD40 repeat-like superfamily protein*), Acc16485.1 (*alpha-glucan phosphorylase*) Acc30767.1 (*UDP-Glycosyltransferase superfamily protein*), Acc08664.1 (*Ammonium transporter*), Acc18987.1 (*MLP-like protein 423*), Acc03527.1 (*AGAMOUS-like*) and Acc08233.1 (*NAD(P) binding protein superfamily*), showed significantly higher expression in Psa-FT and Psa-TMT genotypes compared to Psa-Sus genotypes. Comparatively, genes including Acc01014.1 (*Salycilic acid carboxyl methyltransferase*), Acc24057.1 (*Auxin efflux carrier family protein*), Acc04255.1 (*Acyl-CoA N-acyltransferases (NAT) superfamily*) and Acc13577.1 (*Nudix hydrolase*) were significantly expressed in Psa-Sus compared to Psa-FT and Psa-TMT genotypes.

Further we explored the expression of these genes in leaf tissues of ‘Hort16A’ and P1 plants, inoculated with Psa for bacterial growth assessments (Supplementary Fig. 9), at 0 and 24 hrs time points post-infection. We found that Acc16485.1 (*alpha-glucan phosphorylase*) and Acc03527.1 (*AGAMOUS-like*) were significantly upregulated in P1 at 0 and 24 hrs post-infection compared to ‘Hort16A’ suggesting that their expression is naturally higher in the tolerant parent or suppressed in the susceptible parent and is not induced during early hours of Psa infection (Supplementary Fig. 17). Rest of the candidate genes were not differentially expressed in between the two parents at both time points except Acc08664.1 (*Ammonium transporter*) which was found to be significantly up-regulated in P1 within 24 hrs post-infection compared to ‘Hort16A’ (Supplementary Fig. 17).

## Discussion

This study provides the first information about genetic loci involved in the host-pathogen relationship between *A. chinensis* and Psa. Although a genetic map of the chromosomal location of basal defense and R-genes has been reported ^34^, there has been no previous genetic mapping of Psa resistance. Our study employed natural field and artificial infection data in three environments over multiple years, combined with genetic and transcriptomic experiments in a segregating population resulting from a cross between Psa-susceptible ‘Hort16A’ and a tolerant male P1, to develop an understanding of the genetic factors underpinning quantitative tolerance to Psa in diploid *A. chinensis*.

QTL mapping of the field phenotypic data following natural infection demonstrated the oligogenic nature of this field tolerance, with a single major-effect QTL for tolerance being identified on LG27 in ‘Hort16A’ and six minor-effect QTLs on LGs 3, 14, 15, 22, 24 and 28 of P1. In addition, we demonstrated the interaction of four of the QTLs (LGs 27, 14, 22, 28), accounting for 30 to 40% of the total variance. Our results are consistent with reports of quantitative tolerance against sub-species of *Pseudomonas syringae* ^35–38^ in other hosts and reinforce the long-standing view that no single genetic model can account for incomplete or partial resistance ^39,40^. The major QTL on LG27 of ‘Hort16A’ (initially identified in the field for control of tolerance and expressed as Psa_score_Field) overlaps QTLs for tissue specific responses (Fig. 5). These were for the Ooze phenotype in the stab bioassay in both parents (on LG27.1 S27_4621046 in P1 and LG27 4358305 in ‘Hort16A’) and for the FA_Week3 phenotype in ‘Hort16A’ (on LG27, S27_4853516). In addition, a number of other QTLs identified from the stab bioassay overlapped in the genomic regions S13_6915810 and S13_10678547 on LG13 (Ooze, Tip_death, Stem_necrosis and Psa_score_Stab) (Fig. 5). As stem necrosis leads to collapse of the vascular structure, we suggest that oozing, together with stem necrosis, is not only an important phenotype for assessing tolerance to Psa, but also possibly points towards diverse mechanisms providing field tolerance in *A. chinensis*, that might involve cell wall strengthening and basal defense.

Validation of SSR markers underlying the QTL on LG27 in an independent population of the same cross, as well as another diploid *A. chinensis* population, supports association of this region with Psa tolerance. Genetic analysis of the polymorphism under the LG27 QTL region in ‘Hort16A’ × P1 and other populations indicated that tolerance to Psa is recessive and derived from the male parent of ‘Hort16A’ and that there is likely a susceptibility gene(s) in this region of diploid *A. chinensis*. Further investigation in the kiwifruit germplasm for tolerance-associated haplotypes in this region will aid in fine mapping and the search for candidate gene(s) for Psa tolerance.

Pyramiding of pest and disease resistance loci to enhance durability is an important focus of most crop breeding programs ^40,41^. Marker-Assisted Selection (MAS) has been recognized as a useful tool in breeding perennial fruit crops for major traits such as disease tolerance, flowering, ripening ^42–45^ and is the most efficient route to pyramiding of resistance loci. The first step towards using MAS to improve the efficiency of breeding new Psa-tolerant *A. chinensis* cultivars is the identification of key genetic loci controlling field tolerance to Psa. The moderate-high to high resistance to Psa identified in diploid *A. chinensis* seedlings in PFR breeding populations was reported to be under polygenic control ^25^ and our study has identified a number of genetic loci associated with field tolerance and tissue-specific responses to Psa.

The polygenic nature of tolerance to the pathogen is both an advantage and a disadvantage for breeders. Quantitative resistances that aggregate small effects from multiple genes are relatively durable in comparison to qualitative resistances, as virulent pathovars can more readily evade single *Resistance* (*R*) gene-based resistance ^46,47^. Furthermore, quantitative resistances can also improve the durability of *R*-gene mediated resistances ^48^. However, validation of genetic markers for multiple QTLs in the populations of different ploidy levels that exist in *A. chinensis* can be a challenge. As multiple sources of resistance to *Psa* from a range of species exist in New Zealand kiwifruit germplasm ^25,27,49^, resistance pyramiding based on multiple QTLs is a sustainable first approach in a kiwifruit breeding program and can be strengthened in future with yet unidentified *R* gene resistances against Psa. The oligogenic tolerance to Psa in *A. chinensis* that we have described provides a framework that could lead to the development of durably Psa*-*resistant cultivars.

Pathovars of *P. syringae* have a complex relationship with their hosts ^50^ and develop a range of phenotypes in annual or perennial plant species ^51^. Additional QTLs were identified for associated with tissue-specific responses of *A. chinensis* to Psa in the stab and flood bioassays and some of these overlapped. For example, QTLs for phenotypes in vascular tissues including Stem_necrosis, Stem_collapse and Ooze were adjacent or overlapped on LGs 13 and 16, but QTLs for leaf-associated phenotypes in the stab assay including Wilt, Leaf_spots and Tip death and overall health score recorded in the flood assay (FA_Week1-5) were located on LGs 3, 5, 7, 10 and 18 (Fig 5). This is consistent with a previous finding where distinct quantitative genetic variation underlies leaf and stem specific phenotypic responses to a pathogen ^52^. As the QTLs located using bioassays were not identified for field Psa tolerance, it appears probable that different genetic mechanisms regulate the response to Psa infection in different environments and in different tissues. Many environmental factors differ in greenhouse and in *in vitro* growth conditions compared to the field so might contribute to the plasticity of plant phenotypic responses. This includes factors such as temperature^53–55^, humidity^56–60^, other microbial communities in the field, as well as physiological changes during the growth and aging of *A. chinensis* vines may have an effect. In the future, elucidation of the role of the genetic loci regulating the observed tissue-specific responses to Psa infection will be helpful in determining the dynamics of the host-pathogen relationship in the disease triangle of the *A. chinensis* / Psa patho-system ^61^. Remarkably, a number of the QTLs identified in the bioassays overlie differentially expressed genes, identified from RNA-seq data from field tolerant and susceptible genotypes (Fig. 5).

In general, the association of genes determining quantitative tolerance with a range of mechanisms of innate immunity or PTI enables them to act effectively to counter the virulence strategies of pathogens during different stages of plant development ^62^. In *A. chinensis*, the genome assembly has demonstrated that more genes are associated with PTI, than with *R* gene based Effector-Triggered Immunity (ETI), implying a strong selective pressure on the expansion of genes involved in PTI ^32^. Further evidence for this notion comes from studies exploring the transcriptome of the kiwifruit-Psa interaction the in period directly following inoculation ^63–65^. Data obtained from our study have provided a list of classes of gene families underlying the QTLs that might be directly or indirectly involved in the innate immune response of *Actinidia* and its host-pathogen relationship with Psa over the longer term in the field.

The region underlying the most significant QTL on chromosome 27 are associated with plant defense. (Supplementary Table 2).. A gene encoding a putative cell wall protein Acc15766 (Acc15766.1), located under the P1 LG14 QTL for field tolerance, was employed to design SNP marker E6P3. Two QTLs on LG13 of ‘Hort16A’ were repeatedly identified in association with control of stem necrosis and health, and Psa score in bioassays, as well as in field screens. Underlying these QTLs were two genes, one an orthologue of *Ethylene production protein 1*/*ETO1* (Acc14810.1) that is intricately linked with a plant’s susceptibility to pathogens ^66^, and the other a *Protein ENHANCED DOWNY MILDEW 2*/*EDM2* (Acc14938.1), which is involved in DNA methylation, transcriptional regulation and plant resistance to an oomycete pathogen ^67^.

In the present study we performed RNA-seq on different groups of F1 genotypes from a single population exhibiting extreme variation in field tolerance and susceptibility to natural Psa levels for at least 3 years to explore genes that are associated with Psa tolerance and susceptibility in field over an extended time period. A putative orthologue of *UGT72B1*, which is highly expressed in tolerant Psa_TMT and Psa_FT genotypes and localized within 2-LOD interval of Psa_score_Field QTL on LG 27. Association of *UGT72B1* with non-host resistance against a fungal pathogen has been suggested, as it encodes an enzyme of the phenylpropanoid pathway ^68^. RT-qRT-PCR analysis on samples from controlled inoculation further showed that this gene is induced 24 hrs post-Psa infection in both ‘Hort16A’ and P1, however this needs to be validated if this is the causal gene in Psa tolerance. *EDM2* is significantly upregulated in the field-tolerant Psa-FT genotypes and co-localizes with the QTL on LG13 associated to stem necrosis and collpase. A gene encoding putative cellulose synthase (Acc15562.1), located on the upper arm of LG14, was upregulated in field-tolerant genotypes and might play a role in strengthening the vascular system. On LG24, an orthologue of a Histone protein coding gene (Acc27699.1) that was downregulated in field-tolerant genotypes (Psa-FT and Psa-TMT) underlies a P1 QTL that is associated with field tolerance.

Other gene families that are differentially expressed encode proteins with putative functions associated with PTI, for example detoxification-like protein Acc00747.1 ^69^, a MADS-box like transcription factor Acc03527.1 ^70^, terpene synthases Acc13740.1, Acc13742.1, Acc22685.1, Acc22685.1 ^71^, MLP-like proteins Acc18987.1, Acc13742.1 ^72^ Acc20584.1, Acc20586.1 thioredoxin-like protein ^73^, cellulose synthase-like protein Acc27502.1 ^74^, WD40-repeat containing super-family protein Acc23960.1^75^, UV-B*-*induced protein DUF760 Acc25706.1, Acc14728 ^76^, protein of unknown function (DUF247) /Acc08767, ammonium transporter 2/ Acc08664.1 ^77^. A defense gene that is linked to carbohydrate metabolism that was upregulated in Psa-TMT but downregulated in Psa-FT encodes a putative beta-galactosidase Acc13005.1.

Furthermore, we also verified the expression of the candidate genes associated with plant immunity in Psa-TMT, Psa-FT and Psa-Sus genotypes using gene-specific primers. Consistent with the RNAseq data, we found these genes to be significantly differentially expressed in the tolerant genotypes compared to susceptible genotypes. Specifically, Acc16485.1 (*Alpha-glucan phosphorylase*), Acc03527.1 (*AGAMOUS-like*) and Acc08664.1 (*Ammonium transporter*) genes were confirmed to be significantly induced in P1 in greenhouse and field tolerant genotypes. Acc03527.1 (*AGAMOUS-like*) is located very close to the QTL on LG3 in P1 for Psa_score_Field and an ammonium transporter gene has been recently shown to be involved in stem rust resistance in wheat ^77^. Our study therefore provide new resource for candidate RNA-biomarkers for predicting tolerance in kiwifruit field breeding nurseries that can lead to improve the speed of selective breeding of multi-genic trait ^78^.

Expansion of the pathogenic *P. syringae* strains and their divergence with respect to virulence factors and toxins, as well as antimicrobial compounds ^5,79,80^, indicate that the capabilities of this pathogen in suppressing plant defense are remarkable. Advances in the genomics of both *A. chinensis* and Psa make them a powerful plant–pathogen model system in the context of perennial host species. Results from this study will be utilized to develop MAS for Psa tolerance in diploid breeding populations and to elucidate the molecular mechanisms to fend off the virulent strain of Psa.

## Methods

### Plant material

The populations for genetic mapping of resistance to Psa were progeny of a cross between Psa-susceptible ‘Hort16A’ (female) and tolerant P1 (male) and comprised of three sets of population. The first set, a pilot population comprised 53 genotypes that were clonally propagated 3 to 5 times through cuttings, planted at the PFR Te Puke Research orchard and maintained under standard orchard conditions from 2013 to 2016. The expanded population of 236 ‘Hort16A’ × P1 F1 genotypes was germinated in 2015 aseptically in standard tissue culture growth conditions ^81^. Each genotype was replicated 35–40 times from cuttings, either in tissue culture or under standard greenhouse conditions, prior to field phenotyping or bioassays (Supplementary Fig. 1). Field planting of 230 genotypes (6 to 14 replicates per genotype), was in a randomized block design, in February 2017 at Te Puke and Kerikeri research orchards. A third ‘Hort16A’ × P1 population, of 128 genotypes, was planted in February 2016 in Te Puke and utilized for validation of the genetic markers developed in the other two populations. A diploid *A. chinensis* population ‘Hort22D’ × P2 with 433 individuals was used for marker validation. This population, was maintained under standard orchard conditions from 2011 to 2017 in the Kerikeri research orchard where it was sprayed with antibacterial sprays including copper to suppress disease symptoms, in accordance with the Kiwifruit Vine Health guidelines. We note that the great great grandfather of P2 is the father of ‘Hort16A’.

### Phenotyping

The pilot field population was phenotyped monthly for symptoms from natural Psa infection between 2013 and 2015 and the data used to develop the phenotypic scoring for the expanded population, for which phenotyping was monthly from February 2017 to September 2018. Traits scored included cane death, ooze, shoot death and tip death (Fig. 1). Presence/absence of leaf spots was not recorded, as scores in the pilot study exhibited high between-plant variability. A cumulative Psa score (Psa_score_Field) was calculated, with removals due to oozing and >50% cane death scoring twice as highly as tip death and shoot death. Least squares mean (LS Mean) was calculated for each individual genotype, based on this score.

The bioassays were performed in controlled environments, with the stab bioassay ^26^, being performed between September to December and February to April, in 2016, 2017 and 2018. Inoculations were performed with 10627 SmR, a naturally occurring streptomycin-resistant isolate of Psa biovar 3 ^82,83^, in the greenhouse with temperatures of 22 to 30°C. In total, 200 genotypes were phenotyped using the stab bioassay, with 35 batches phenotyped across three years. Details are in Supplemental Methods S1. The flood bioassay ^29^ was performed by flooding six biological replicates of each genotype with Psa, that had been grown on tissue-culture media in an aseptic growth medium in a tub for 4 to 6 weeks. Details are provided in Supplemental Methods S1.

### Bacterial inoculations for assessment of growth curve in resistant vs. susceptible plants

Assessment of the growth curve for Psa in ‘Hort16A’ and P1 was performed using multiple biological replicates in the greenhouse, as described for the stab test bioassay. Young potted kiwifruit plants were inoculated with Psa, on 8 to 10 biological replicates of each genotype in February, 2018. Further details are provided in Supplemental Methods S1.

### Genotyping, genetic maps and QTL mapping

DNA was extracted from freeze-dried leaves using the Cetyl trimethylammonium bromide (CTAB) method ^84^. GBS libraries were prepared for 53 individuals from the pilot population and 236 individuals from the expanded population, as well as the two parents, using the method described in detail by ^85^, modified from the standard GBS protocol ^30^. The individual and pooled libraries were checked for quality with a Fragment Analyser (Advanced Analytical) and pooled libraries with satisfactory QC were dried down and dispatched to the Australian Genome Research Facility (AGRF) for single-end sequencing on an Illumina® HiSeq™ platform. The sequencing reads were de-multiplexed based on GBS library preparation barcodes using the eautils.1.1.2-537 package and those reads starting with the approved barcode immediately followed by the remnant of the *Bam*HI cut site sequence were retained for further analysis. Variant calling and genotyping was performed using TASSEL v3.0 and 5.0 and ~60,000 and 80,000 SNP calls were generated for the individuals in the two populations, respectively. SNP calling was performed using an early version of the Red5 genome (1.68.5), which preceded the 1.69.0 version ^33^, and the ‘Hongyang’ genome ^32^ as references. Genome coordinates for the Red5 version 1.68.5 were converted to those of the published version using in-house PERL scripts (available on request from Ross Crowhurst, PFR). The coordinates for SNPs associated with the QTL peaks in the published Red5 genome are listed in Supplementary Data1. The SNP calls represented 70% coverage of the expanded population. In our data sets (Supplementary Data 1), Red5 markers begin with S, whereas markers generated from ‘Hongyang’ begin with HY, followed by the number of the linkage group and the position of the marker on the respective physical genome (for example S1_10661198 or HY10_1385907). The SNP data were subsequently filtered to obtain 9,875 and 9,327 SNP markers polymorphic between ‘Hort16A’ and P1, respectively (3,364 for P1 in pilot study). JoinMap v 5.0 ^86^ was used to develop genetic linkage maps for both the parents, at a LOD score between 15 and 22. QTL mapping was performed using the rQTL package ^87^ and MapQTL5 software ^88^. Multiple QTL models, including Maximum likelihood (EM), Haley-Knott regression, Non-parametric, multiple imputation and Kruskal-Wallis analysis (KW) were employed for single QTL scans.

### RNA-seq and RT-qRT-PCR

Total RNA was extracted from healthy young leaves, at the sixth to ninth position from the apical leaf, from genotypes in the field that were segregated into three groups based on susceptibility to Psa. The first two groups were of two-year-old plants that were defined respectively as: 1) tolerant/medium tolerant (Psa-TMT), consisting of three relatively Psa-tolerant genotypes, including P1, and 2) susceptible (Psa-Sus), comprising three fully Psa-susceptible genotypes including ‘Hort16A’ based on observations of 8 to 11 biological replicates. The third group (Psa-FT) exhibited field tolerance to Psa over four years. RNA extraction was from samples snap frozen with liquid nitrogen, using the Spectrum Total Plant RNA kit (Sigma-Aldridge, Auckland, New Zealand) and QC was performed with the Fragment Analyzer to select RNA with RNA Integrity Number (RIN) of 7.1-8.2. Samples from three biological replicates were pooled. Library preparation at the Australian Genome Research Facility used the TruSeq Stranded kit and subsequent paired-end Illumina® sequencing employed the NovaSeq6000 platform. An average of ~19 million, 150 bp paired-end reads were retrieved for each sample (~6 Gb) and read sequences of low-quality, ribosomal RNA as well as adaptors were filtered out using Trimmomatic ^89^ and SortMeRna ^90^. RNA-seq reads were aligned to the Red5 reference gene models using STAR and differential expression analysis was performed using DESeq2 ^91^. The details concerning the target file, read statistics and DEGs are provided in Supplementary Data2.

For RT-qRT-PCR on field samples, RNA was extracted (as described above) from 3-7 clonal replicates of each genotype from Psa-TMT and Psa-Sus group from heathy leaf tissues. RNA was extracted from leaf tissues of 5 different genotypes in Psa-FT group, as these genotypes are not clonally replicated. For RT-qRT-PCR on infected tissues, samples were harvested from the leaf tissues used for assessment of bacterial growth curve as described above. A 10mm leaf disc was harvested from the region infected with Psa, at 0, 6, 24 and 48 hrs post-infection, and frozen in liquid nitrogen. One leaf disc was harvested from a single clonal replicate of ‘Hort16A’ and P1 per time point and three biological replicates were harvested at each time point. Data from 0 and 24 hrs time is only presented in the study. Total RNA (~ 2 μg) was treated with DNase I (Roche Applied Sciences) and used for cDNA synthesis using SuperScript IV Reverse Transcriptase (Life Technologies-Invitrogen). The cDNA was diluted 20-fold and used for qRT-PCR employing a LightCycler^®^ 480 SYBR Green 1 Master PCR labelling kit (Roche Applied Sciences) and RotorGene 3000 Real time PCR machine (Corbett Research, Sydney, Australia). Relative transcript abundance was determined relative to the mean of the expression of *Actin and Ubiquitin* genes in the same sample. Comparative quantification was performed as described ^92^. Primers used for genes are provided in Supplementary Table 6.

### SSR and SNP marker design and screening

Repeats were identified manually in the genome sequence underlying the QTLs. PCR primers for SSR markers were designed using Primer3 and employed to screen DNA extracted from the populations ^93^. Analysis and scoring of the alleles in the amplicons was performed on a Hitachi ABI3500 Applied Biosystems genetic analyzer. Primers were also designed around SNPs in the genes identified in the genomic sequence of Red5 underlying the QTLs. The SNP markers were screened using real-time High Resolution Melting analysis ^94^. All primer sequences are provided in Supplementary Supplementary Table 6

## Acknowledgements

We would like to acknowledge AgMARDT New Zealand for a post-doctoral fellowship to JT and funding the work. Financial support was also provided by; 1) PFR Strategic Science Investment Funds Breeding Technology Development, 2) Kiwifruit Breeding Programme and 3) KRIP Phenotying Bioassays. We acknowledge the assistance of Belinda Diepenheim, Andrew Mullan, Renata Blissett, Bruce Dobson, Matt Spier, Judith Rees, Janet Phipps, Deidre Cornish, Janet Yu, Jenny Oldham and Jaqui Wallace.

## Author Contributions

JT performed pathogen infection experiments, DNA extractions, GBS, RNA-seq, RT-qRT-PCRs, data analysis, genetic map construction, QTL mapping, designed the SNP and qPCR markers. JT, SEG and DC wrote the manuscript. LG performed field phenotyping. SH performed stab assay. HB developed SSR markers and performed validation of the markers. CB, AC, KT and MM performed flood assay. EM, AW and KF performed replication of the genotypes in the tissue culture and greenhouse. CW performed validation of the breeding parents from the germplasm. CD performed GBS analysis. RC developed circos view of QTLs and RNA-seq data. MK performed DNA extraction for the validation population. JT, DH and LG performed statistical analysis of the data. JV performed bacterial growth assessments. JM performed validation of the LG27 QTL in the breeding germplasm. KH managed orchard plantations of the genotypes.

---

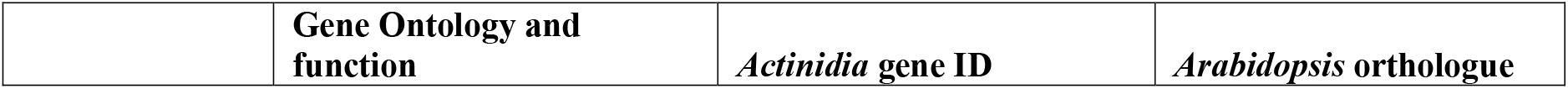

