## Supplemental Methods S1 for "Multiple quantitative trait loci contribute tolerance to bacterial canker incited by *Pseudomonas syringae* pv. *actinidiae* in kiwifruit (*Actinidia chinensis*)"

*Phenotyping*

a. *Stab bioassay*

Each genotype was replicated 8 to 12 times and 3 to 6 biological replicates of each genotype were distributed over 2 to 3 batches of 20 to 50 cm tall potted plants. Psa inoculum was prepared by suspending bacterial cells in sterile reverse osmosis (RO) filtered water to make a concentrated stock (~1-4 x 10^9^ Colony-forming unit (cfu)/ml). This was adjusted to 2 x10^8^ cfu/ml in Phosphate Buffered Saline (PBS) using optical density (OD) readings from a spectrophotometer (NovoSpec, Amersham Biosciences, Auckland, New Zealand) and a standard curve developed using a range of concentrations of Psa checked by measuring and by determining cfu/ml by plating on KB medium. Single inoculations were performed on each plant with a mechanical stab device, consisting of a double row of needles spaced 1.5 mm apart, dipped in the Psa inoculum, on the elongating supple region of the stem, between the top and the middle leaves. Each batch included ‘Hort16A’ as a susceptible control, with *A. arguta*, *A. chinensis* var. *deliciosa* and *A. chinensis* var. *chinensis* (*2x* and *4x*) genotypes, supplementing P1 as tolerant controls, as clonal propagation of P1 was difficult. Phenotypes were recorded at the third week post-infection included plant height, Stem_necrosis, Stem_collapse, Leaf_spots, Wilt, Tip_death and Ooze (Fig. 1). A total Psa score (Psa_score_Stab) was calculated using the equation

Psa_score = {Collapse score (Tip_death*2+Stem collapse*5+Total collapse*12) + Stem score (IF Stem_necrosis > 100 score is = 3, IF Stem_necrosis > 10 score is = 2, IF Stem_necrosis > 3 score is = 1) + Ooze + Leaf_spot + Wilt + Stunted tip score*0.5 + Stem break score*0.5 + Regrowth score*0.5 + Callus/wound score *0.5}.

Mock inoculations performed with 10mM MgCl_2_ resulted in no stem lesions > 0.7 cm, or other Psa phenotypic responses in ‘Hort16A’ or F1 genotypes evaluated.

Phenotypic data were analysed using appropriate statistical models. For Stem_necrosis as a proportion of plant height and for Psa_score_Stab, linear mixed models were fitted (REML procedure in GenStat, version 17, VSNi Ltd, Hemel Hempstead, UK, 2014), with random effects for inoculation batch, genotype and inoculation batch x genotype. Means for each genotype were predicted from the model. For Ooze, Leaf_spots, Wilt, Tip_death and Stem_collapse, which were recorded as present or absent for each plant, Bernoulli generalized linear mixed models were fitted (GLMM procedure in GenStat v17), with random effects for inoculation batch, genotype and inoculation batch x genotype. The residual deviance was fixed as 1. Means for each genotype were predicted from the model on the logit scale and back-transformed to obtain predicted percentages. Repeatability of clonal means was calculated following ^1^, dividing σ^2^bc by 3.5 (the average number of blocks a line was present in) and σ^2^w by 3.5 x 1.9 (the average number of replicates of a line within a block).

b. *Flood assay*

Psa inoculum was prepared by re-suspending bacterial cells in 10mM MgCl_2_ at ~1 x10^7^ cfu/ml, with a final concentration of 0.005% Silwet L-77. The flooding completely covered the plants for 3 mins. The inoculum was then drained from the tub and the same inoculum was used to flood six additional tubs. This bioassay was performed from 2016 to 2018 under controlled conditions of 16hrs light/8hrs dark at 20°C as the plants became available. Each genotype was grown in triplicate in the tub and two tubs of each genotype were inoculated in the same batch. Phenotypes were recorded individually on a scale of 0 to 4, with 0 completely healthy and 4 sick or dead, weekly for five weeks post-infection. An average score was calculated for each genotype per time point using the equation

Average health score per time point (FA_Week) = (n*0)+(n*1)+(n*2)+(n*3)+(n*4) where n = number of replicates of the genotype scored for the respective stage.

Mock inoculations performed by flooding ‘Hort16A’ and other F1 genotypes with 10mM MgCl_2_ resulted in no adverse phenotypes.

*Bacterial inoculations for assessment of growth curve in resistant vs. susceptible plants*

Inoculum was prepared freshly from agar plates of King’s B medium ^2^ on which Psa had grown for 48 hours at 25°C. Bacteria were re-suspended in sterile water with 0.03% Du-Wett® (Etec Crop Solution, Auckland New Zealand) to a final concentration of 2 x10^8^ cfu/ml. Four leaves per plant were marked with five areas on each side of the main vein. Each area on one side of the leaf was inoculated with 20 µl of the bacterial suspension, the other side received 20 µl of sterile water. The oldest and youngest leaves were not used. All plants were then placed into high-humidity tents (RH>90%) for a week, after which the relative humidity was reduced to about 50%, by opening two 20 x 20 cm flaps that had been cut into the top of the humidity tents. Epiphytic and endophytic populations of Psa were determined two hours, 3 days and 10 days post inoculation. At each time point, one disc (7mm diam.) from one inoculated area and one water-treated control area were sampled per leaf. In total there were three inoculated discs and three control discs per plant per time point. The epiphytic population of Psa was determined by washing the leaf discs individually in 1ml of sterile water. The washings were serially diluted 1:10 and three 10 µl drops of each dilution were plated on King’s B media supplemented with streptomycin (100mg/L). The numbers of colonies were counted following two days’ incubation at 28°C. The endophytic population of Psa was determined from the washed discs. The discs were surface sterilized by washing them individually in 1% sodium hypochlorite for 3 mins, rinsing in sterile water, washing in 70% ethanol for 3 mins, rinsing in sterile water, washing in 1% sodium hypochlorite for 3 mins followed by three rinses in sterile water. Three 20 µl drops of the final rinse were plated on King’s B plates without antibiotic to check that all epiphytic bacteria had been killed. The discs were then transferred to 2 ml vials containing 5 ceramic beads and 300 µl of sterile water. Samples were macerated in a FastPrep2 (MP Biomedicals) for 6.0 m/s for 40s. Tubes were centrifuged for 2mins in a Gyrozen centrifuge to pellet plant material, before removing 100 µl of supernatant, which was serially diluted and three 10 µl drops of each dilution plated onto King’s B medium supplemented with streptomycin (100 mg/L). When necessary, the identity of the strains growing on King’s B supplemented with streptomycin was confirmed by PCR or duplex PCR as previously described ^3,4^. The experiment was performed on 8 to 10 biological replicates of each genotype in February, 2018.
