## Supplementary Figures and Tables for "Multiple quantitative trait loci contribute tolerance to bacterial canker incited by *Pseudomonas syringae* pv. *actinidiae* in kiwifruit (*Actinidia chinensis*)"

a)

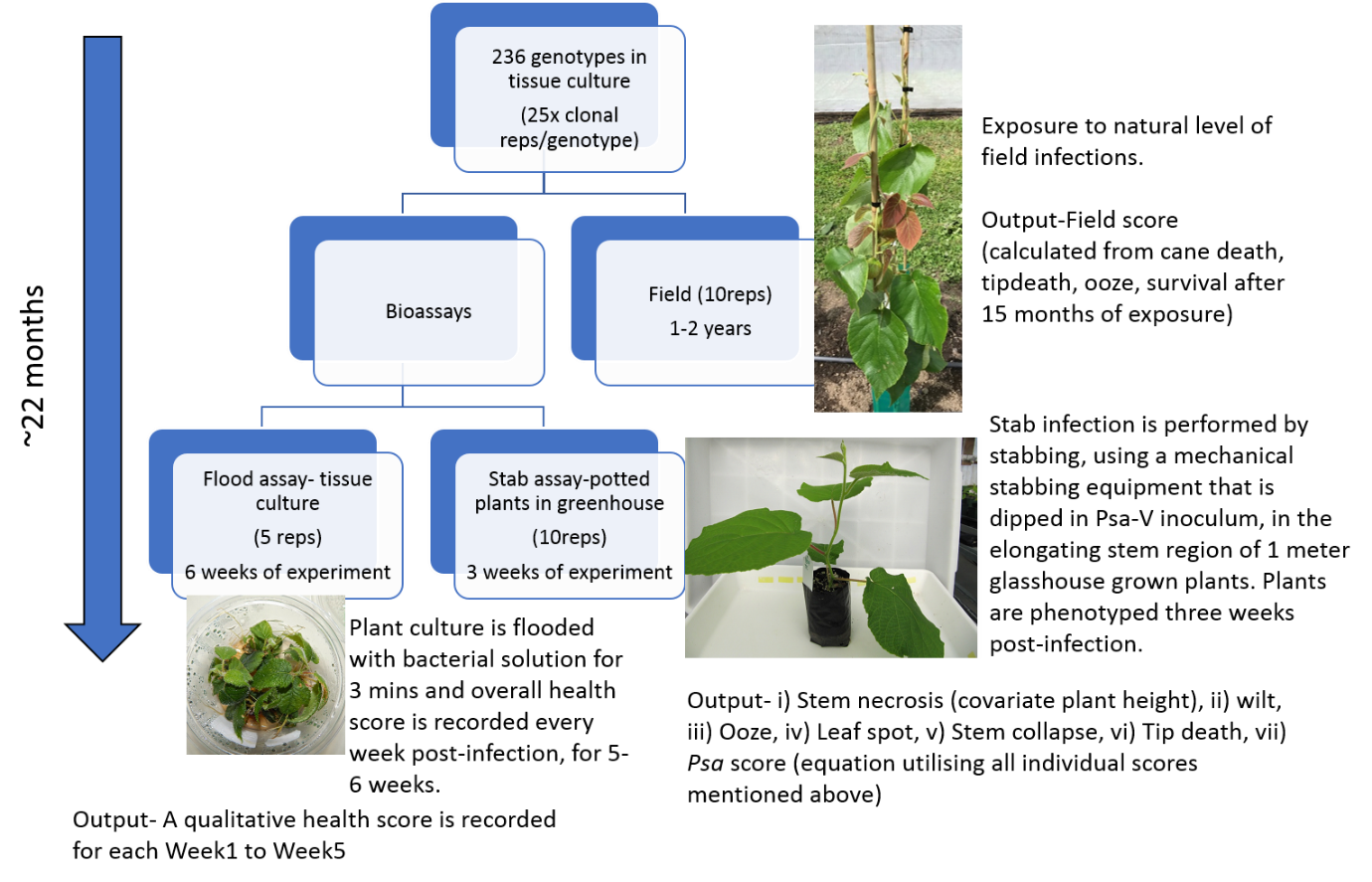

b)

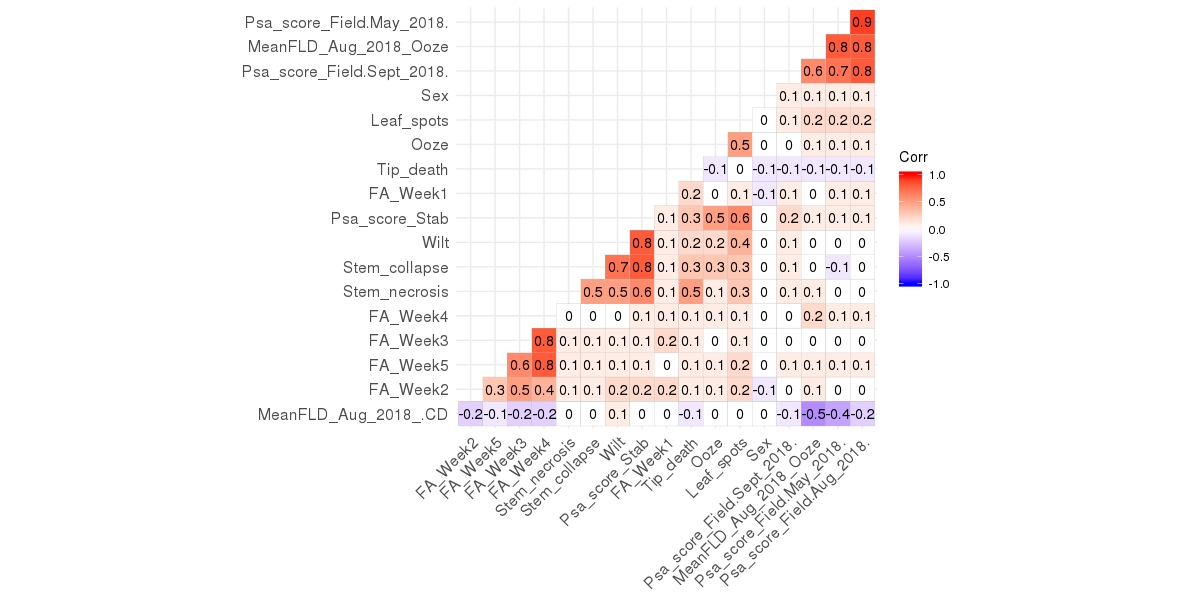

**Supplementary Figure 1**. Workflow for intensive phenotyping of the expanded ‘Hort16A’xP1 population using bioassays and a field trial and correlation between the phenotypes. A) Clonal replicates of genotypes for two bioassays and the field trial were phenotyped during a 22 month time period. B) Correlation between all the phenotypes recorded during field assessments (Psa_score_Field_* and MeanFLD_*), stab assay ( Leaf_spots, Ooze, Tip_death, Psa_score_Stab, Wilt, Stem_necrosis and Stem_collapse) and flood assay (FA_Week*). From 236 genotypes, that were genotyped and phenotyped, 177 genotypes were common in all the phenotypic assessments. Correlation coefficient over 0.15 has a correlation significantly (p<0.05) higher than would be expected purely by chance (the threshold is 0.19 or over for p<0.01, and 0.24 or over for p<0.001).

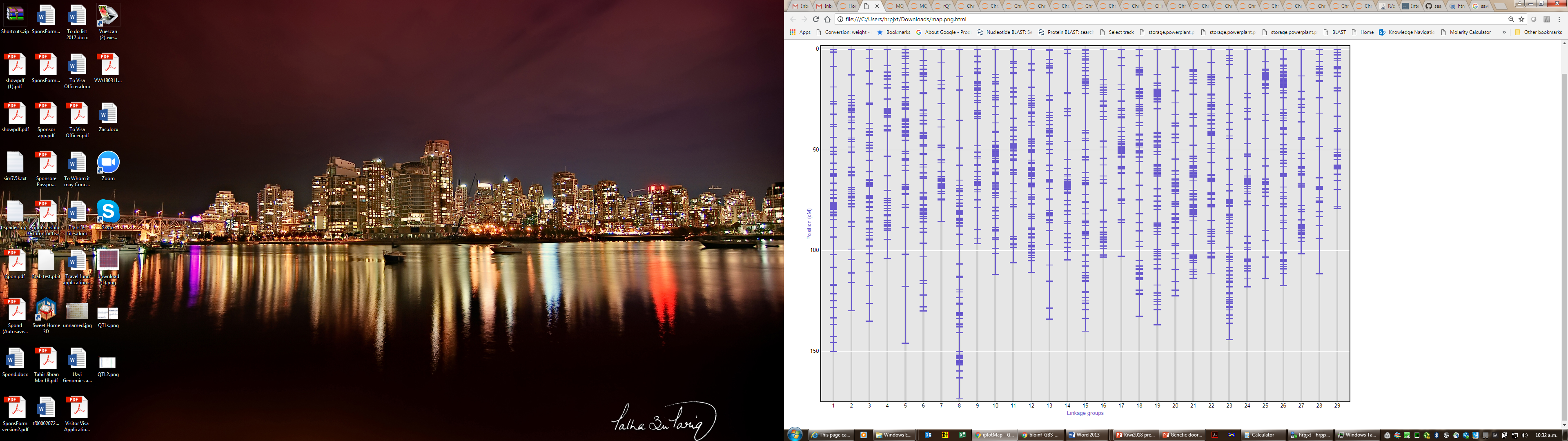

**Supplementary Figure 2**. Genetic maps of ‘Hort16A’ linkage groups in the expanded population.

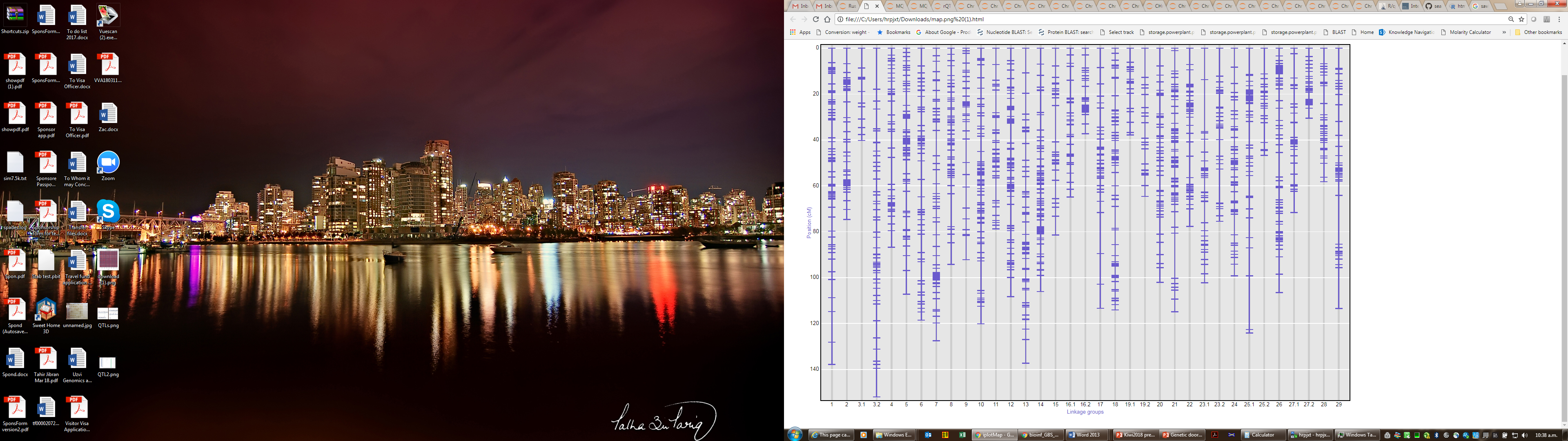

**Supplementary Figure 3**. Genetic maps of P1 linkage groups in the expanded population.

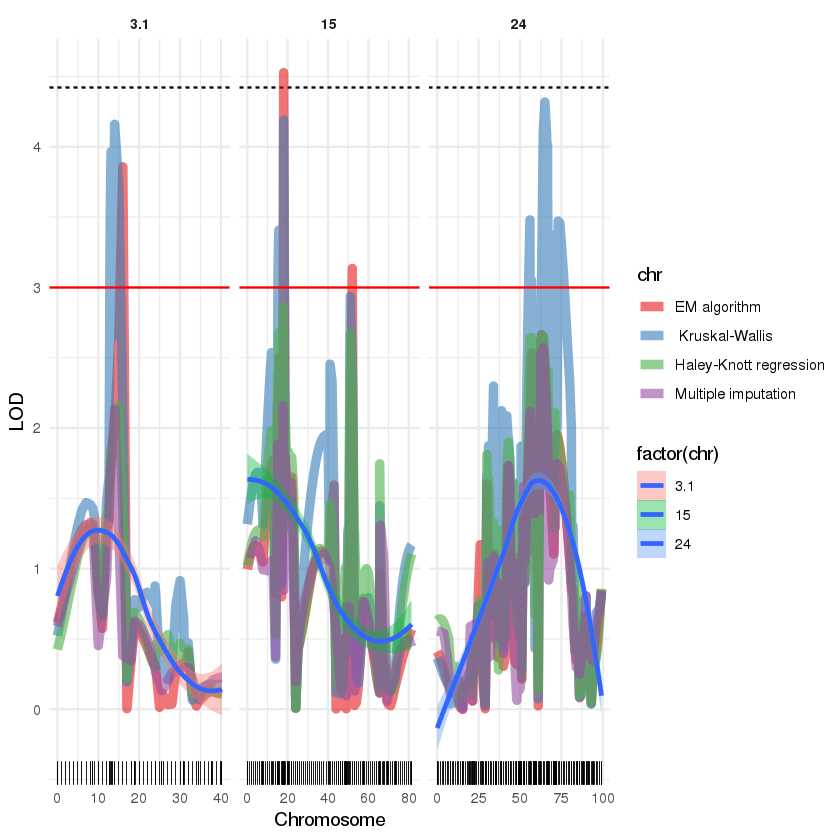

S15_7775365

S24_12069300

S3_1039842

cM

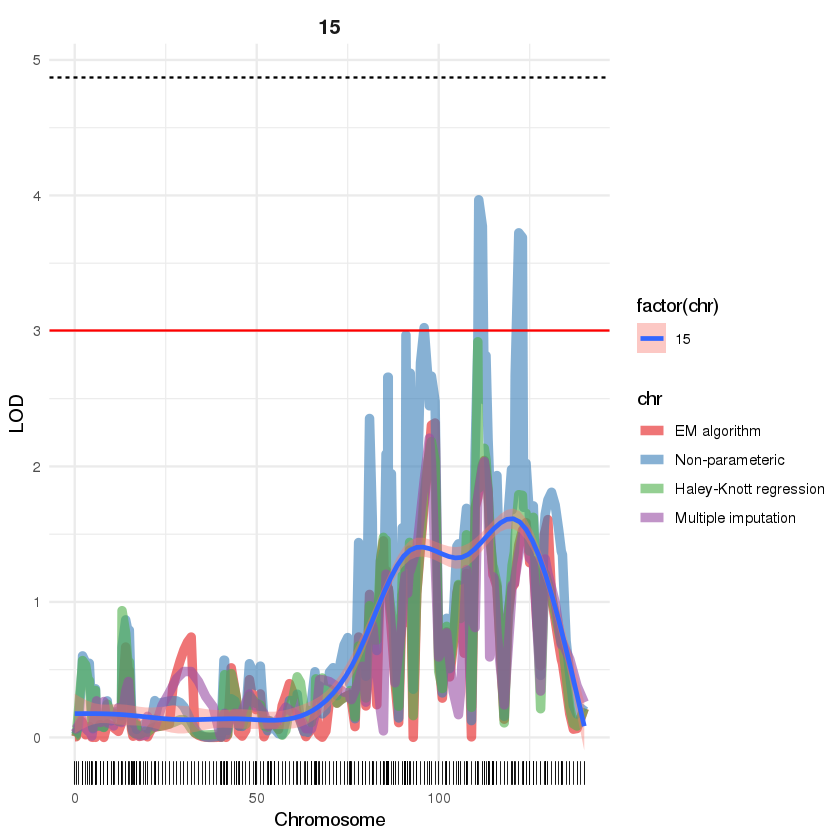

**Supplementary Figure 4**. QTLs in P1 of LG, 3.1, 15 and 24, using various models, for control of the Psa_score_Field phenotype. Black bars represent cM position in the linkage maps for GBS markers. A red line is at LOD3 and dotted line represents the threshold of significance of the QTL at 95 % genome-wide permutation test.

**SSRLG28_1378F5R5**

‘Hort16A’ =st X P1 =rs

**LG14 E6P3(SNP) Acc15766**

Tol

Sus

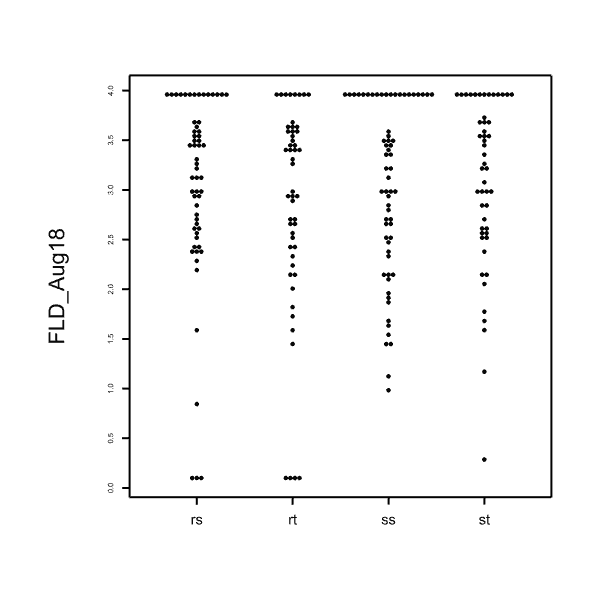

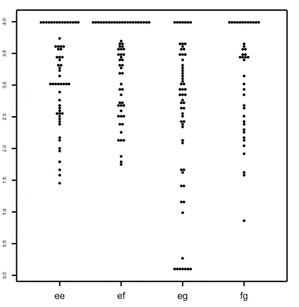

‘Hort16A’ =e**f** X P1 =e**g**

rs

rt

st

ss

***fg***

*ee*

*ef*

*eg*

**Supplementary Figure 5**. Dot plot analysis of effect of alleles of a SNP and an SSR marker underlying the LG14 and LG28 QTLs and the Psa_score_Field phenotype determined in the expanded population after two years in the field.

a) ‘Hort16A’xP1

SSRLG27_4396125F4R4

‘Hort16A’ = *uv* x	P1 = *wx*, where the favourable allele is *v* = 428bp

|  | alive | dead(4) |  |
| --- | --- | --- | --- |
| *wv* & *xv* | 27 | 30 | *wv* & *xv* |
| *wu* & *xu* | 10 | 60 | *wu* & *xu* |

b) ‘Hort22D’ X ‘P2’

SSRLG27_4396125F3R3

‘Hort22D’ = ab x ‘P2’ =aa , where the favourable allele is a = 424 bp

|  | alive | dead(4) |
| --- | --- | --- |
| *ab* | 96 | 78 |
| *aa* | 139 | 59 |

**Supplementary Figure 6**. Association of LG27 haplotypes with Psa tolerance. a) shows segregation of the v allele 428 bp, amplified using SSRLG27_4396125F4R4 marker, for dead and alive vines in a one year old ‘Hort16A’xP1 population. b) shows segregation of the a allele 424 bp, amplified using SSRLG27_4396125F3R3 marker, for dead and alive vines in a five year old population of an *Actinidia chinensis* diploid population from a cross between ‘Hort22D’ and ’P2’ parents.

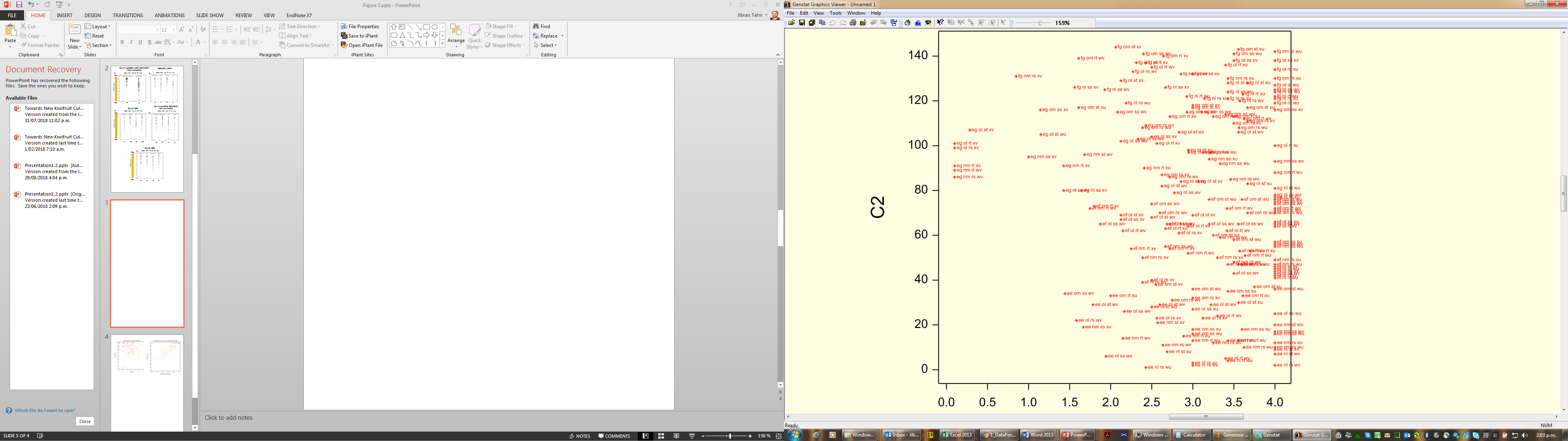

Psa_score_Field

Tolerant

Susceptible

Combination of alleles from 4 QTLs

**Supplementary Figure 7**. Combination of alleles from markers underlying 4 quantitative trait loci (QTLs), LG27, LG22, LG14, LG28, plotted against Psa_score_Field. The parental allelotypes exhibited by the markers for these QTLs are: LG27, ‘Hort16A’ exhibits *uv* and P1 *wx*, with the favourable allele being *v*; LG22, ‘Hort16A’ exhibits *lm* and P1 *no*, with the favourable allele combinations being *nm* and *ol*; LG14, ‘Hort16A’ exhibits *ef* and P1 *eg,* with the favourable allele being *g*; LG28, has *st* from ‘Hort16A’ exhibits *st* and P1 *rs*, with the favourable allele combinations being *rs* and *rt*. The following combinations of alleles show association with tolerance: *eg nm rs wv*, *eg nm rt wv*, *eg nm rt xv*, *eg ol rs xv, eg ol rt xv, eg ol st xv, fg nm rs xv* and *eg nm ss xv*.

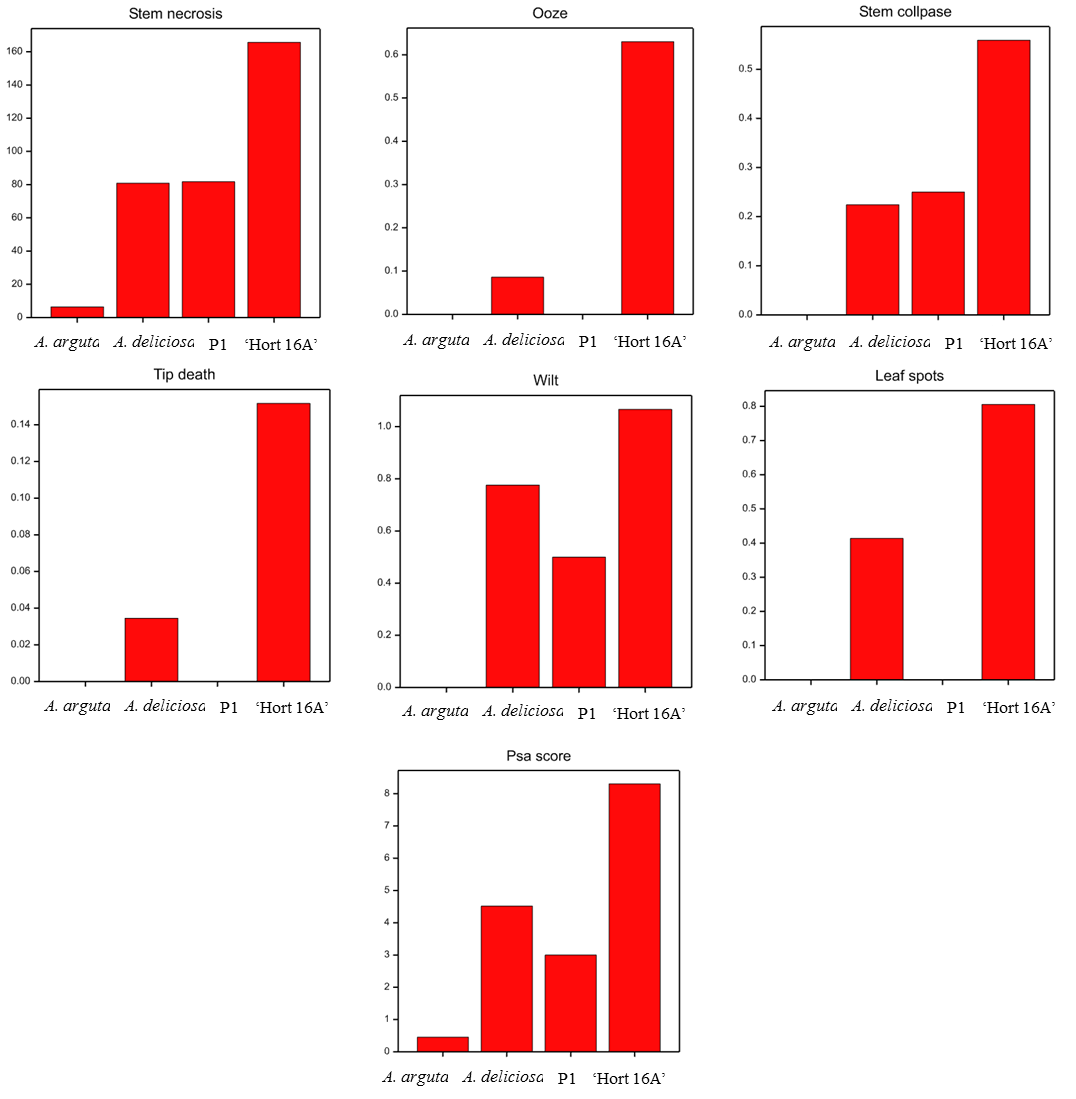

Psa_score_Stab

Leaf_spots

Wilt

Stem_collapse

Ooze

Tip_death

Stem_necrosis

**Supplementary Figure 8**. Means of phenotypic scores in the stab bioassay for ‘Hort16A’, P1,*Actinidia* *arguta* and *A. deliciosa*.

a) b)

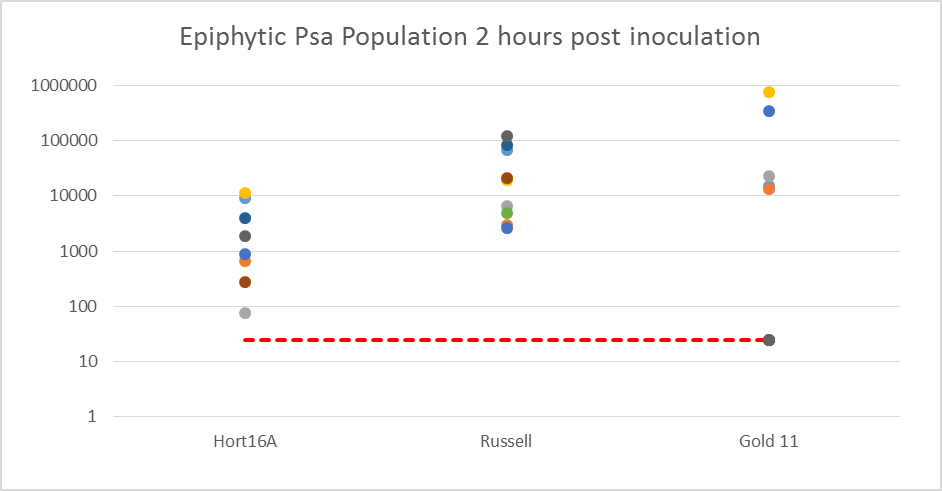

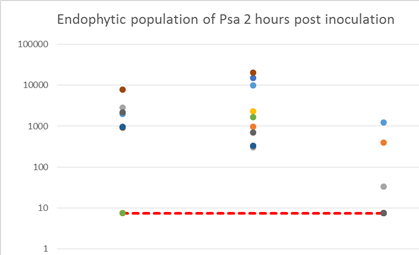

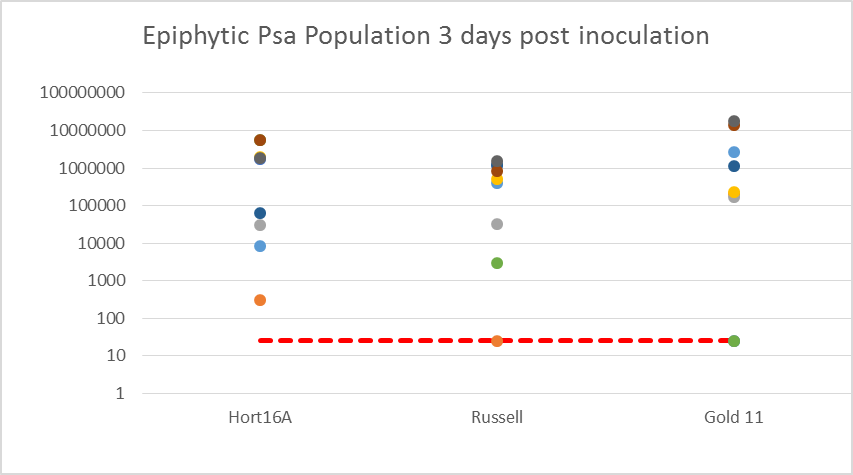

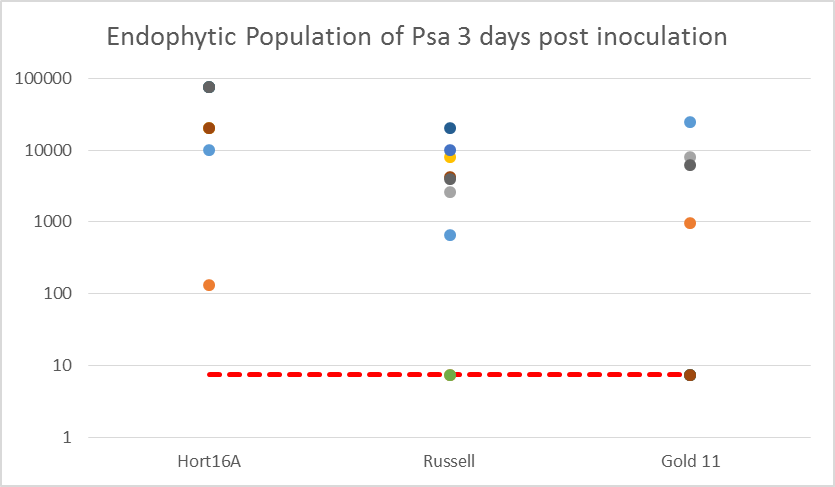

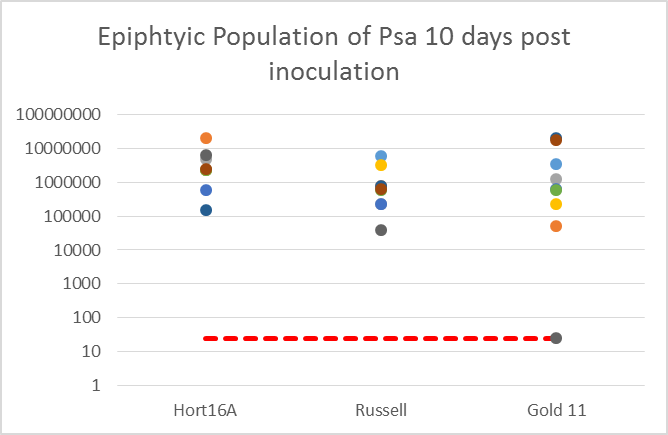

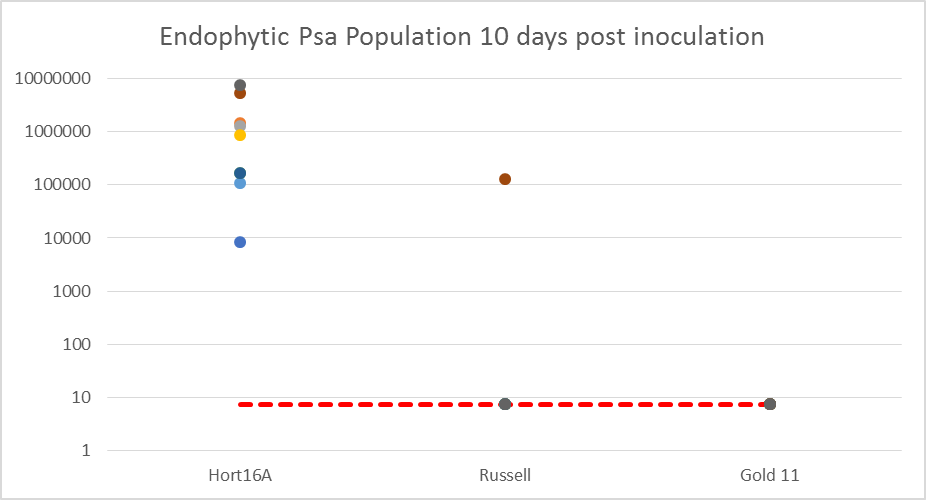

‘Hort 16A’	P1	*4x A. chinensis*	‘Hort 16A’	P1	*4x A. chinensis*

**Supplementary Figure 9**. Epiphytic (a) and endophytic (b) levels of *Psa Biovar 3* in ‘Hort16A’, P1 and a *Psa Biovar 3* tolerant tetraploid (4x) *Actinidia chinensis* genotype at three time points post-infection. The figure depicts the epiphytic and endophytic population levels (colony-forming-unit/cfu) of *Psa Biovar 3* on the Y-axis, at different time points post-inoculation in leaf discs, with eight biological replicates, represented by different coloured circles, for each genotype. Inoculations were performed with 20 ul of a bacterial suspension of *Psa Biovar 3* strain SmR123 containing 2 x 10^8^ cfu/ml. The red broken line indicates minimum threshold detectable for colony counts.

a) Stem_necrosis	b) Ooze

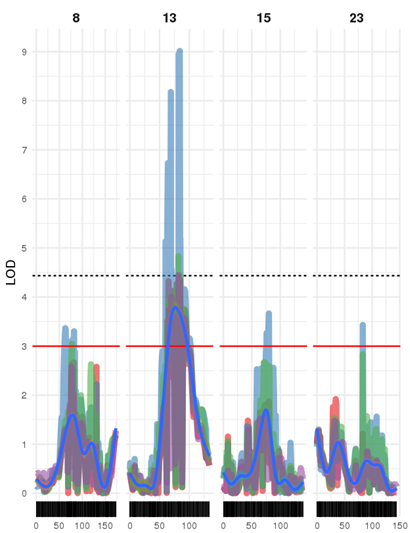

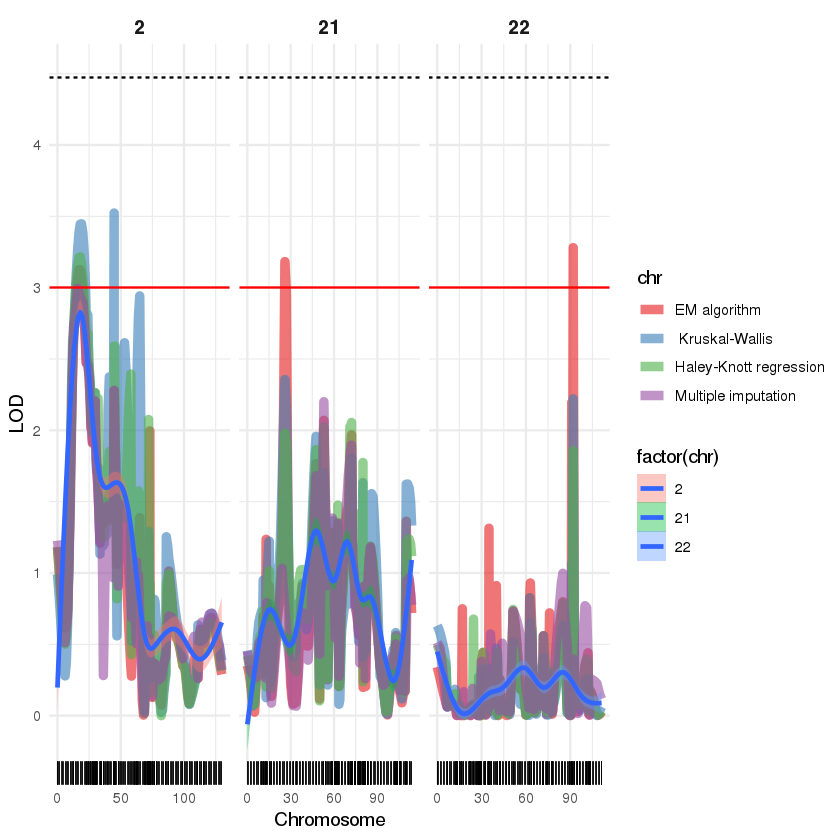

c22.loc92

c21.loc26

c23.loc83

S15_11279533

S13_10678547

S13_6915810

S13_6476279

c8.loc64

S2_4100973

c) Leaf_spots	d) Wilt

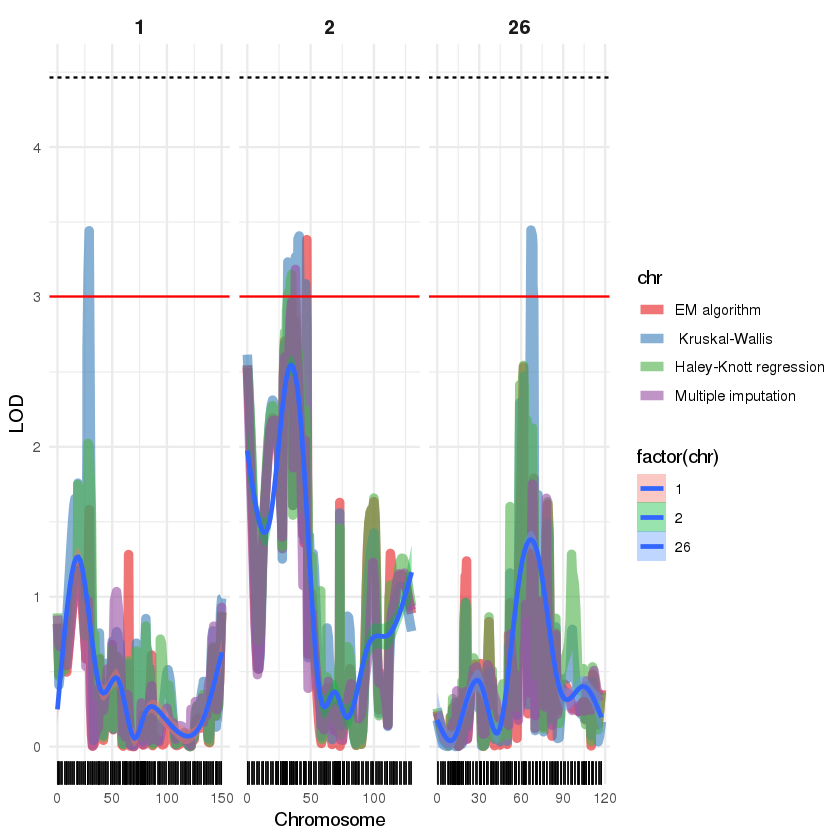

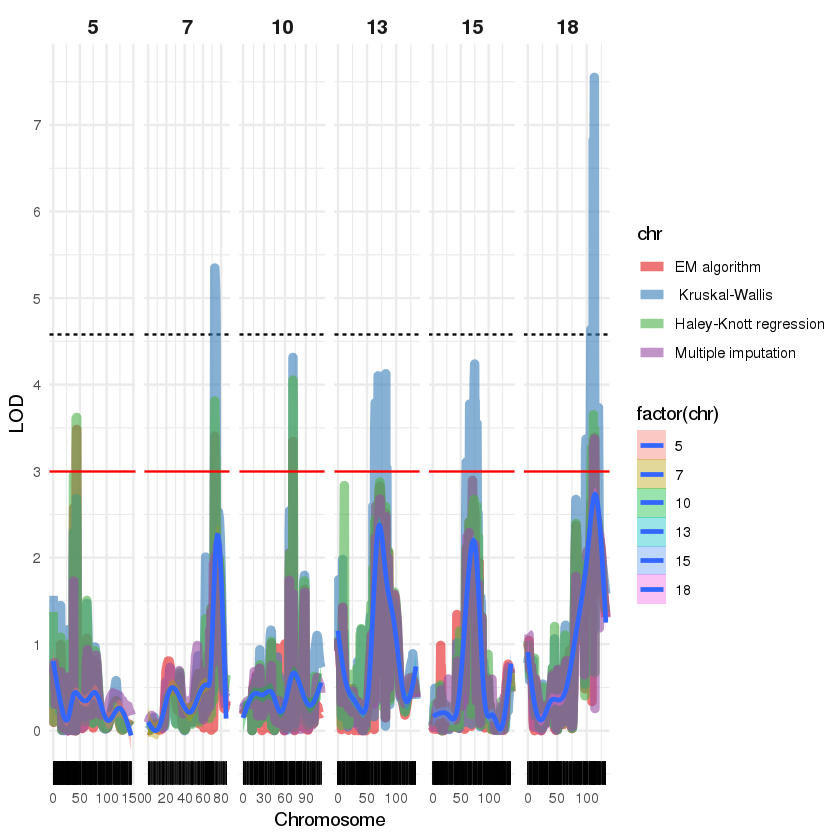

S18_19172999

c15.loc74

S13_10678547

S13_6915810

HY30_1328264

S17_14039379

S26_14412151

c2.loc33

c1.loc29

e) Stem_collapse	f) Psa_score_Stab

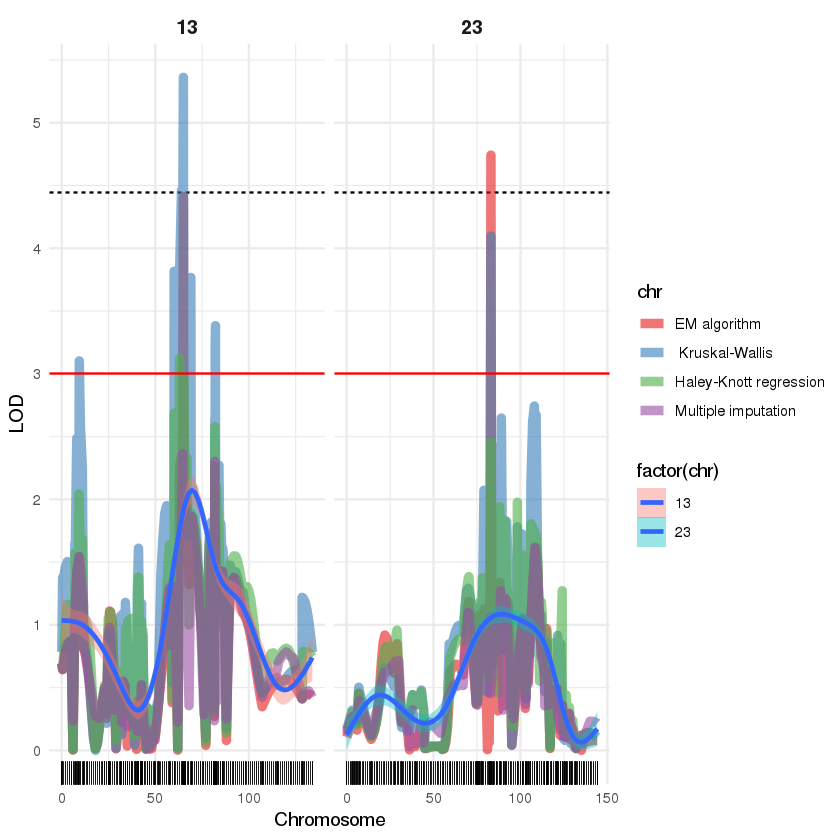

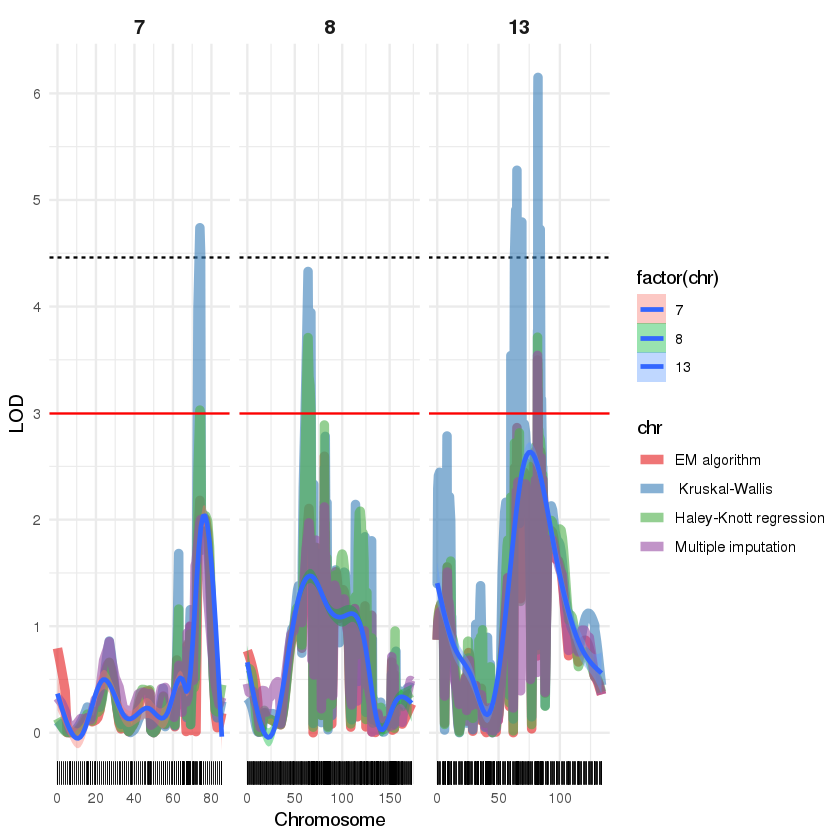

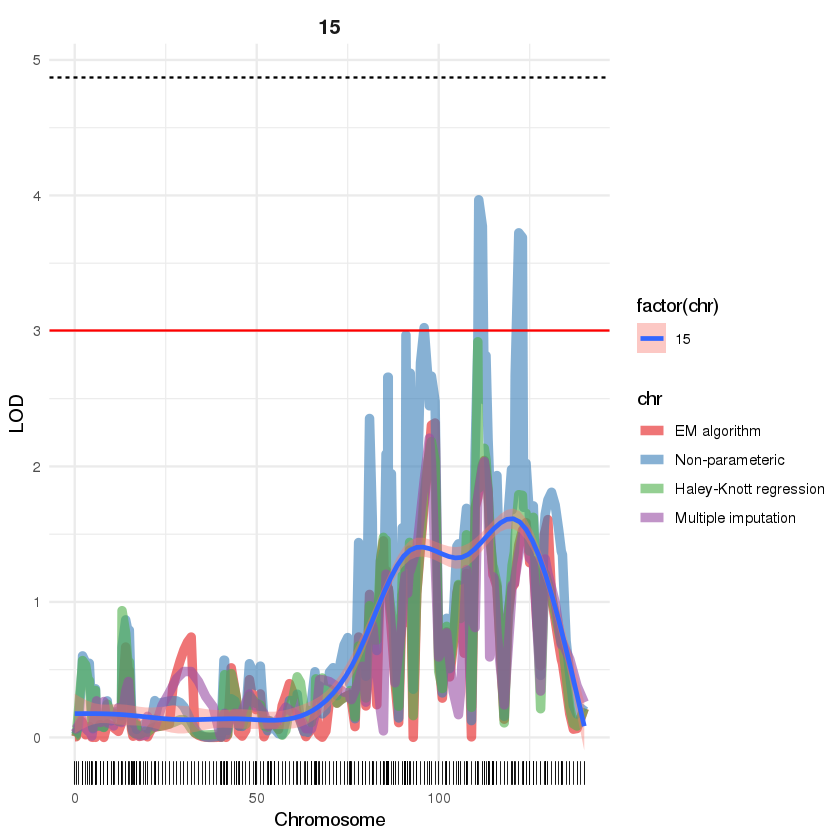

c23.loc83

S13_10678547

c8.loc64

S17_14039379

c13.loc65

S13_6476279

**Supplementary Figure 10**. QTLs on ‘Hort16A’ linkage groups using various models for interval mapping and different phenotypes (a-f) in the Stab bioassay. Black bars represent cM position in the linkage maps for genotyping by sequencing (GBS) markers. A red line indicates LOD 3 and the dotted line the threshold of significance of the QTL at 95 % in a genome-wide permutation test. SNPs with maximum association are indicated for each peak whereas peaks lacking an underlying marker are identified with the predicted genetic position.

a) Stem_necrosis	b) Ooze	c) Leaf_spots

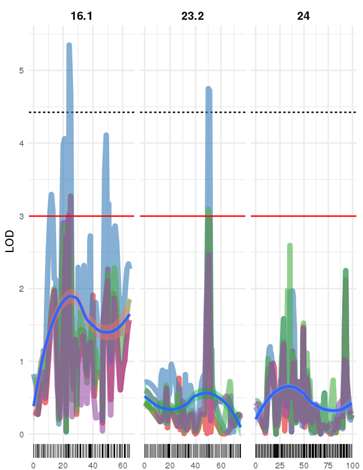

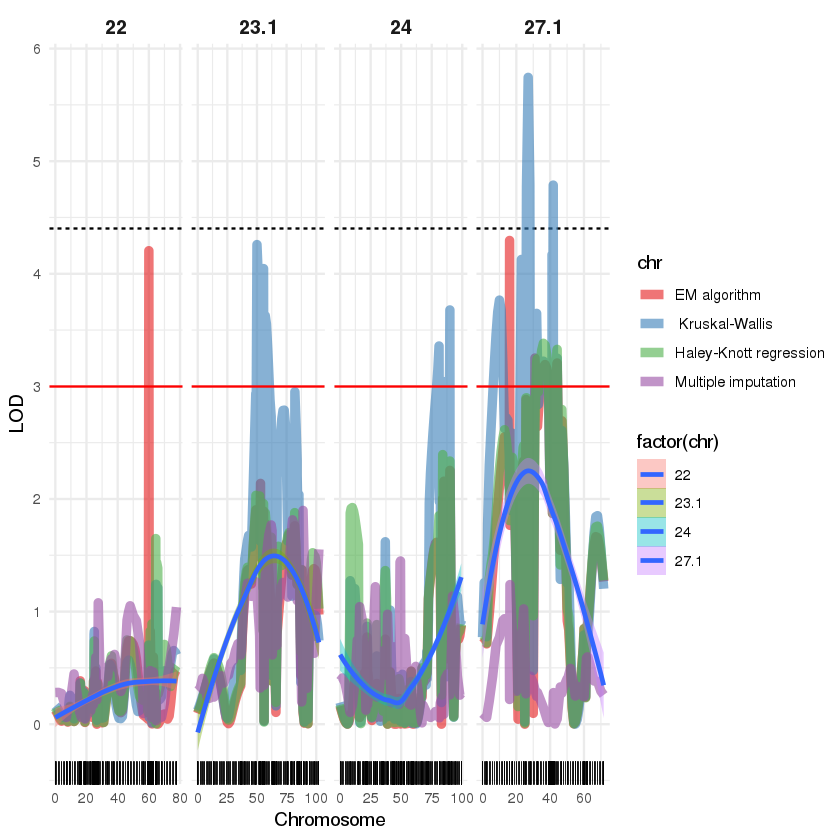

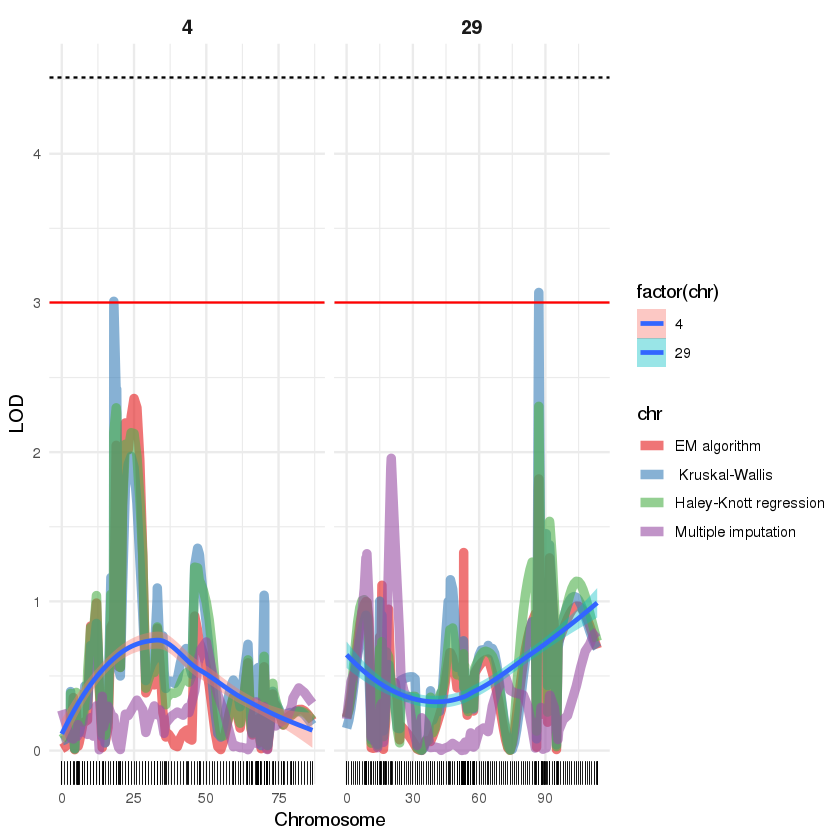

c29.loc87

c4.loc18

c27.1.loc27

S27_3476163

S27_4396089

c24.loc81

c23.1.loc50

c22.loc60

S23_17796340

S16_2518074

c16.1.loc24

d) Wilt	e) Stem_collapse

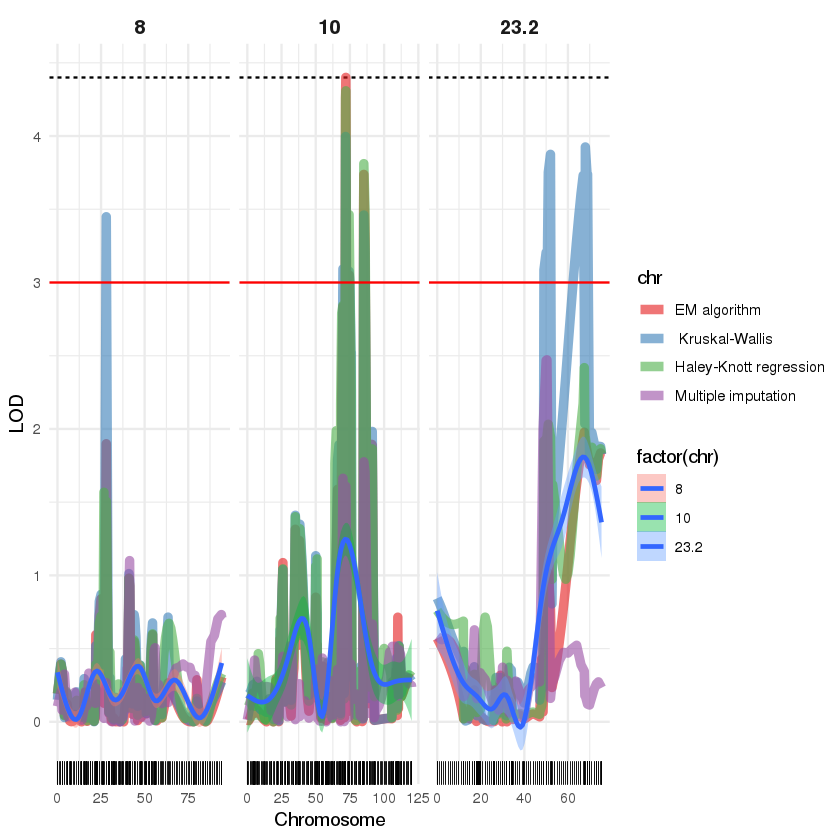

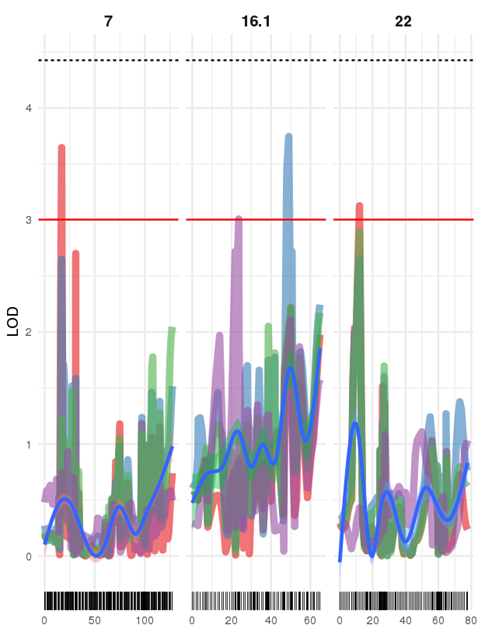

c22.loc12

c16.1.loc20

HY7_17317578

S23_19369514

HY30_1618642

c8.loc28

f) Psa_score_Stab

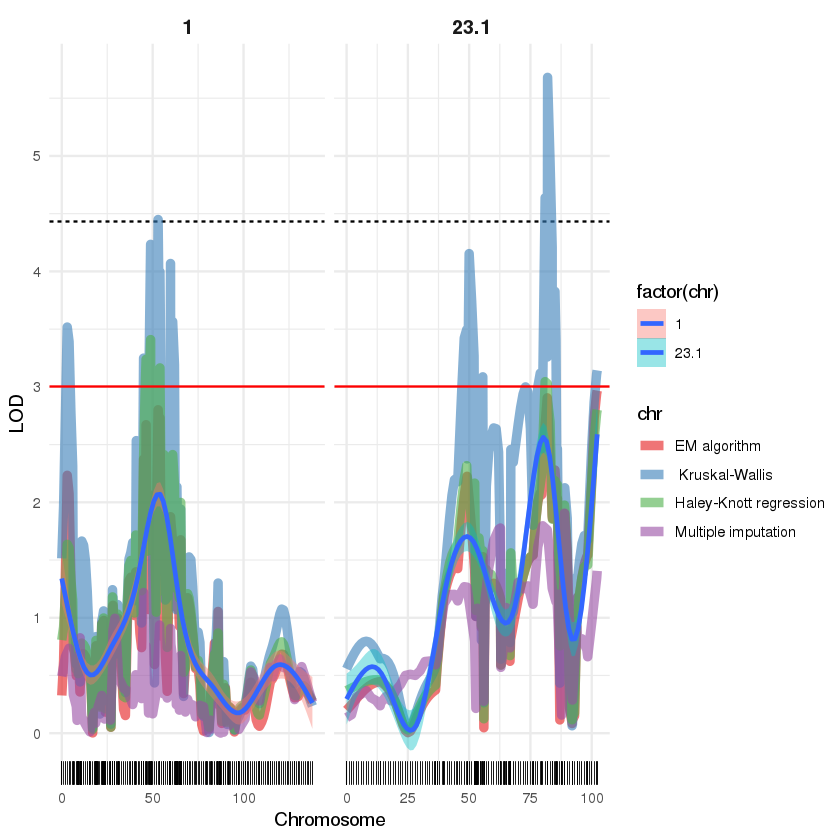

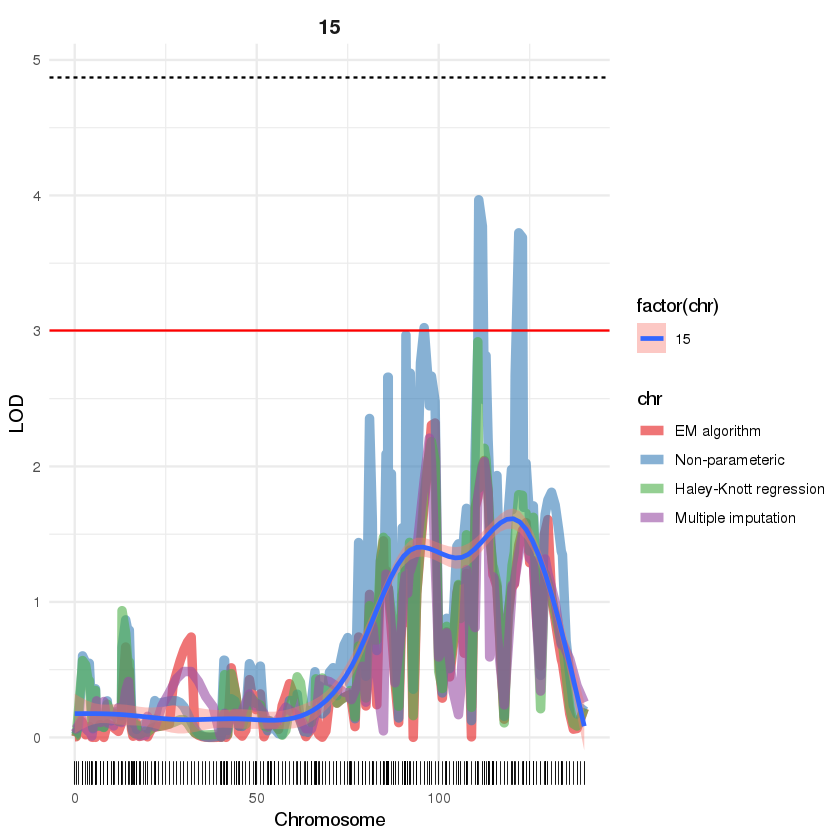

c23.1.loc82

S23_6065543

c1.loc53

S1_8216009

**Supplementary Figure 11**. Quantitative trait loci (QTLs) on P1 linkage groups using various models for interval mapping and different phenotypes (a-f) in the Stab bioassay. Black bars represent cM position in the linkage maps for genotyping by sequencing (GBS) markers. A red line indicates LOD 3 and the dotted line the threshold of significance of the QTL at 95 % in a genome-wide permutation test. SNPs with maximum association are indicated for each peak whereas peaks lacking an underlying marker are identified with the predicted genetic position.

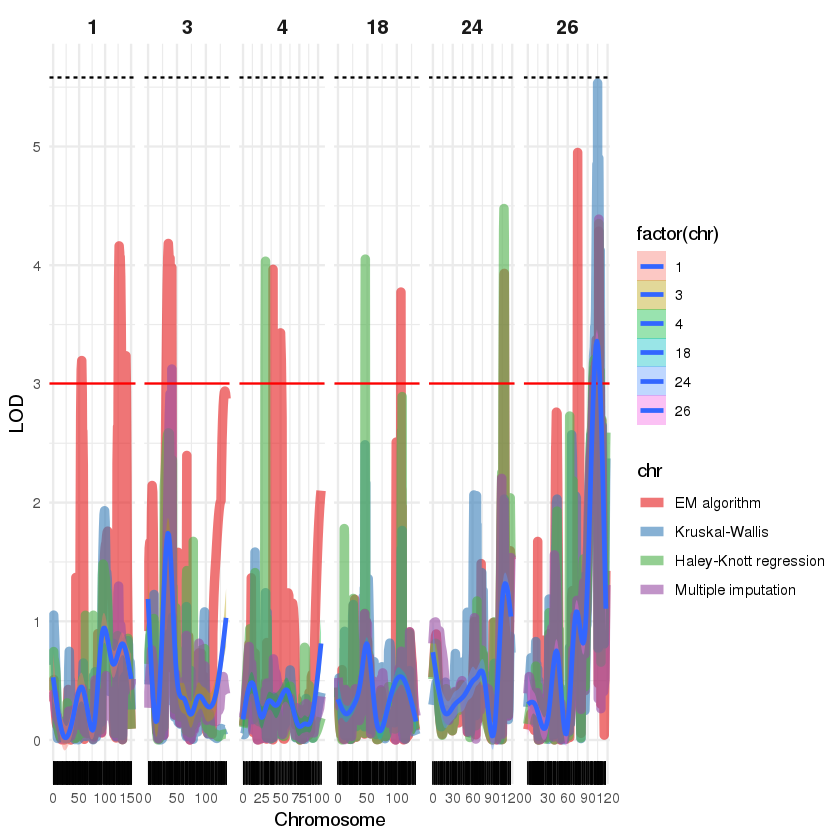

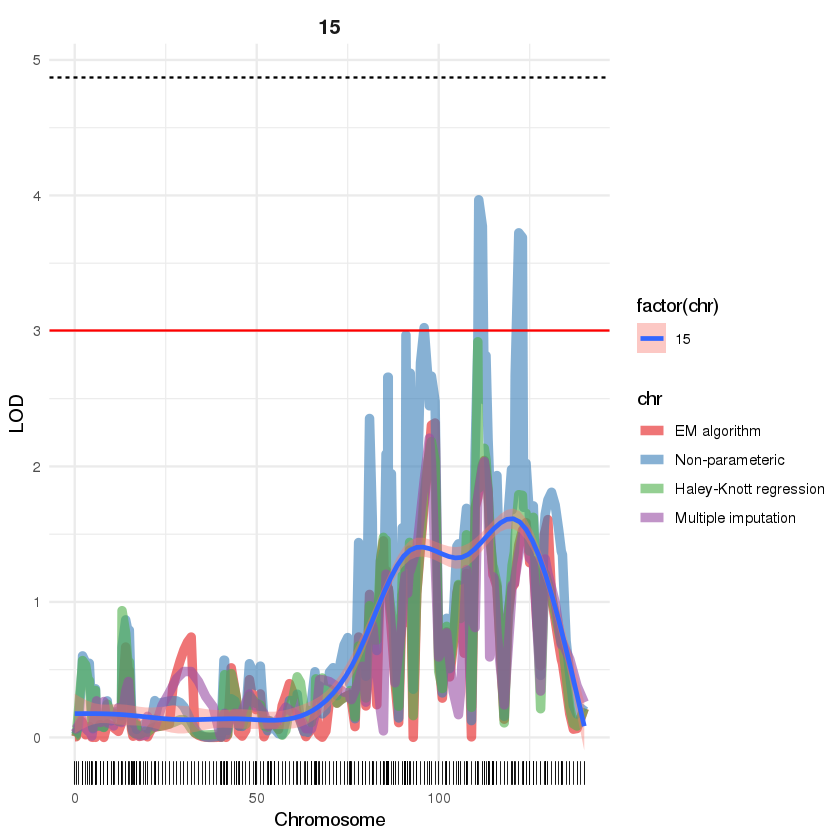

**Supplementary Figure 12**. Quantitative trait loci (QTLs) on ‘Hort16A’ linkage groups using various models for interval mapping for Tip_death phenotype in the stab bioassay. Black bars represent cM position in the linkage maps for genotyping by sequencing (GBS) markers. A red line indicates LOD 3 and a dotted line for the threshold of significance of the QTL at 95 % in a genome-wide permutation test.

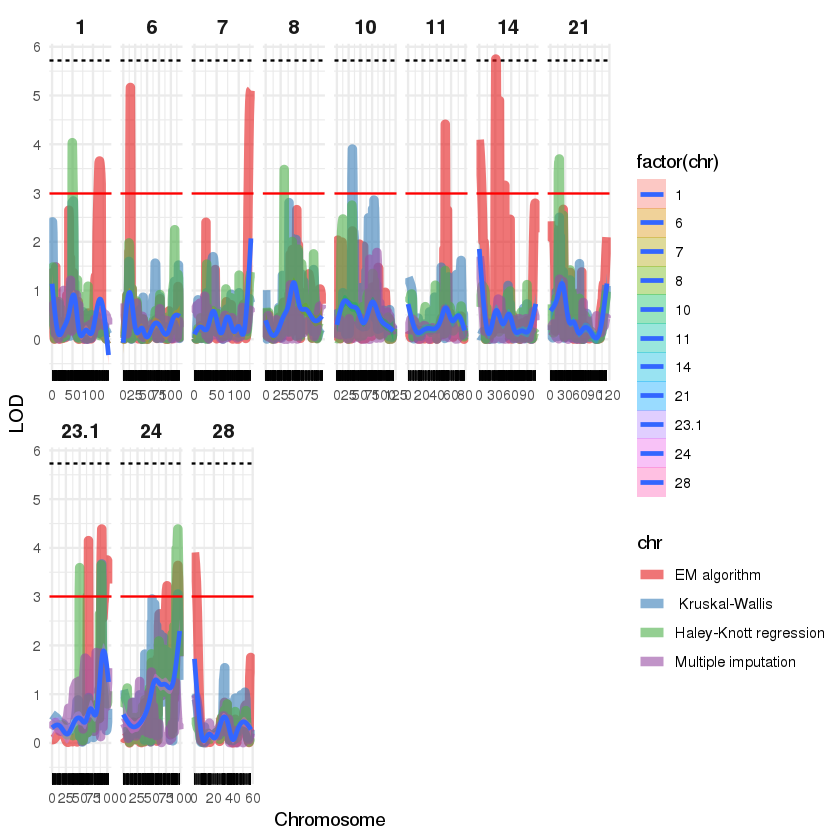

S14_2394123

S28_5625626

S10_3720973

S8_4688024

HY7_17317615

HY21_11458399

S24_14459417

S23_8830490

HY11_9463505

S13_15166616

S8_2653784

S1_17369059

**Supplementary Figure 13**. Quantitative trait loci (QTLs) on P1 linkage groups using various models for interval mapping and Tip_death phenotype in the Stab bioassay. Black bars represent cM position in the linkage maps for genotyping by sequencing (GBS) markers. A red line indicates LOD 3 and the dotted line the threshold of significance of the QTL at 95 % in a genome-wide permutation test. SNPs with maximum association are indicated for each peak whereas peaks lacking an underlying marker are identified with the predicted genetic position

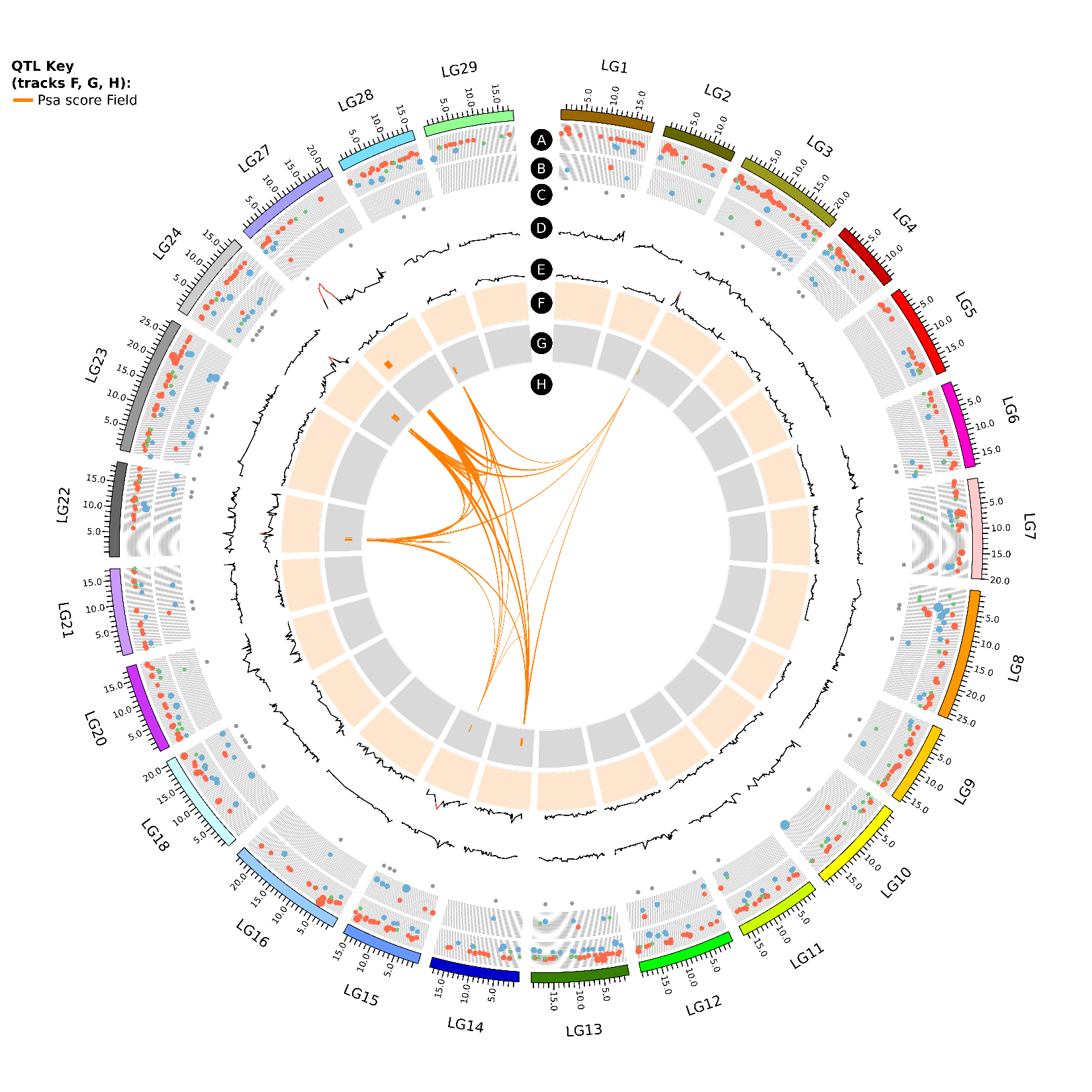

**Supplementary Figure 14**. Circos plot of quantitative trait loci (QTLs) for Psa_score_Field phenotype as well as RNA-seq data associated with *Psa* tolerant and susceptible genotypes, anchored on the chromosomes of the Red5 genome version 1.6.9 associated with QTLs. Tracks A and B represent differentially expressed gene (DEGs) with logFC ±2 and above in fully tolerant (Psa-FT) vs susceptible (Psa-Sus) and tolerant to medium tolerant (PsaTMT) vs Psa-Sus genotypes, respectively. A) blue circles are upregulated and red circles are downregulated genes in Psa-FT compared to Psa-Sus genotypes. B) blue circles are downregulated and red circles are upregulated genes in Psa-TMT compared to Psa-Sus genotypes. Genes with logFC between -1 and 1 are represented by green circles. Increase in circle diameter indicates increasing logFC value. C) DEGs common to Psa-FT and Psa-TMT. Track D and E are LOD values for Psa_score_Field for ‘Hort16A’ and P1 respectively. The lines change from black to red for a LOD score >3.Track F and G are QTLs detected from the phenotype listed in QTL key  in ‘Hort16A’ and P1 respectively. H represents the lines connecting QTLs for similar phenotype on different chromosomes.

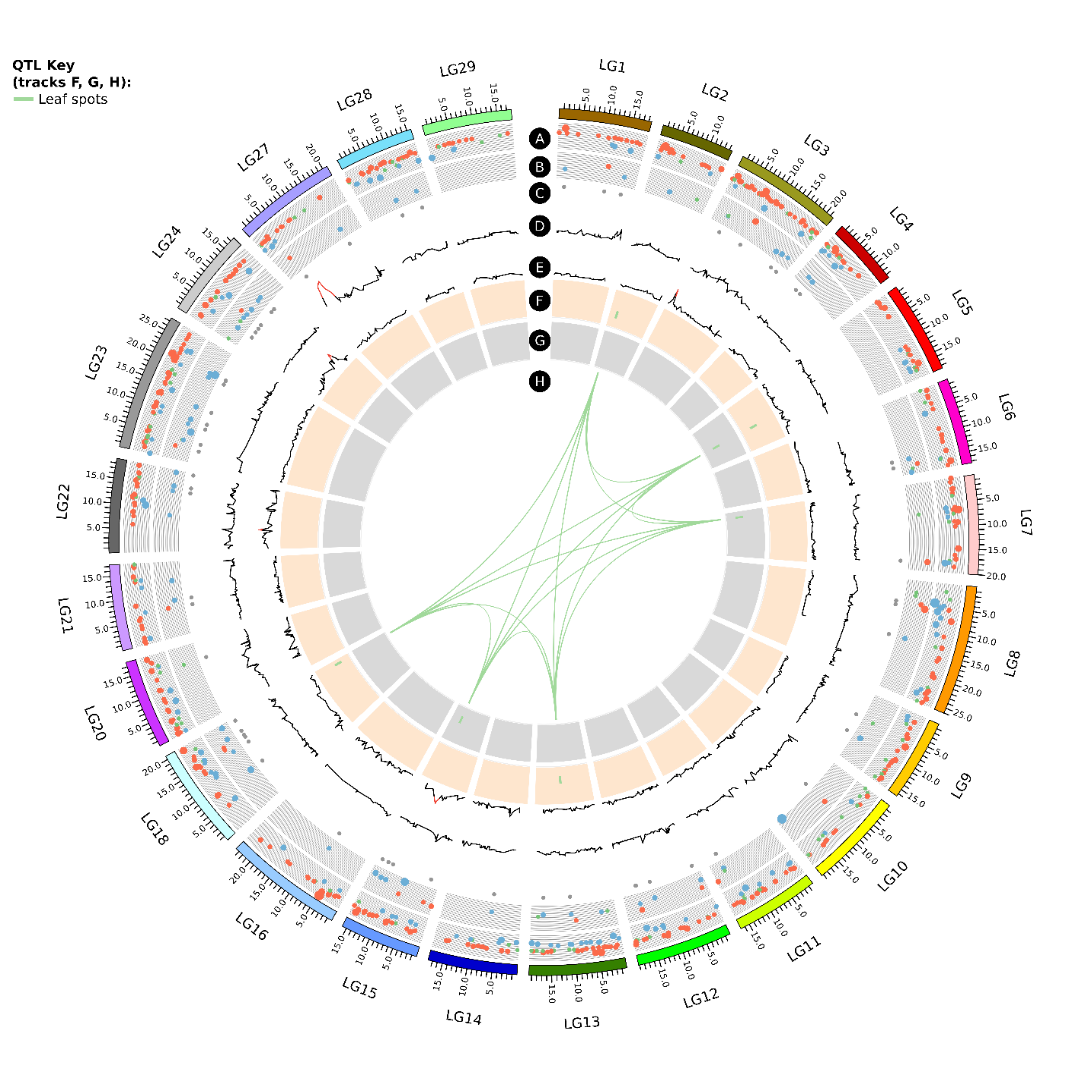

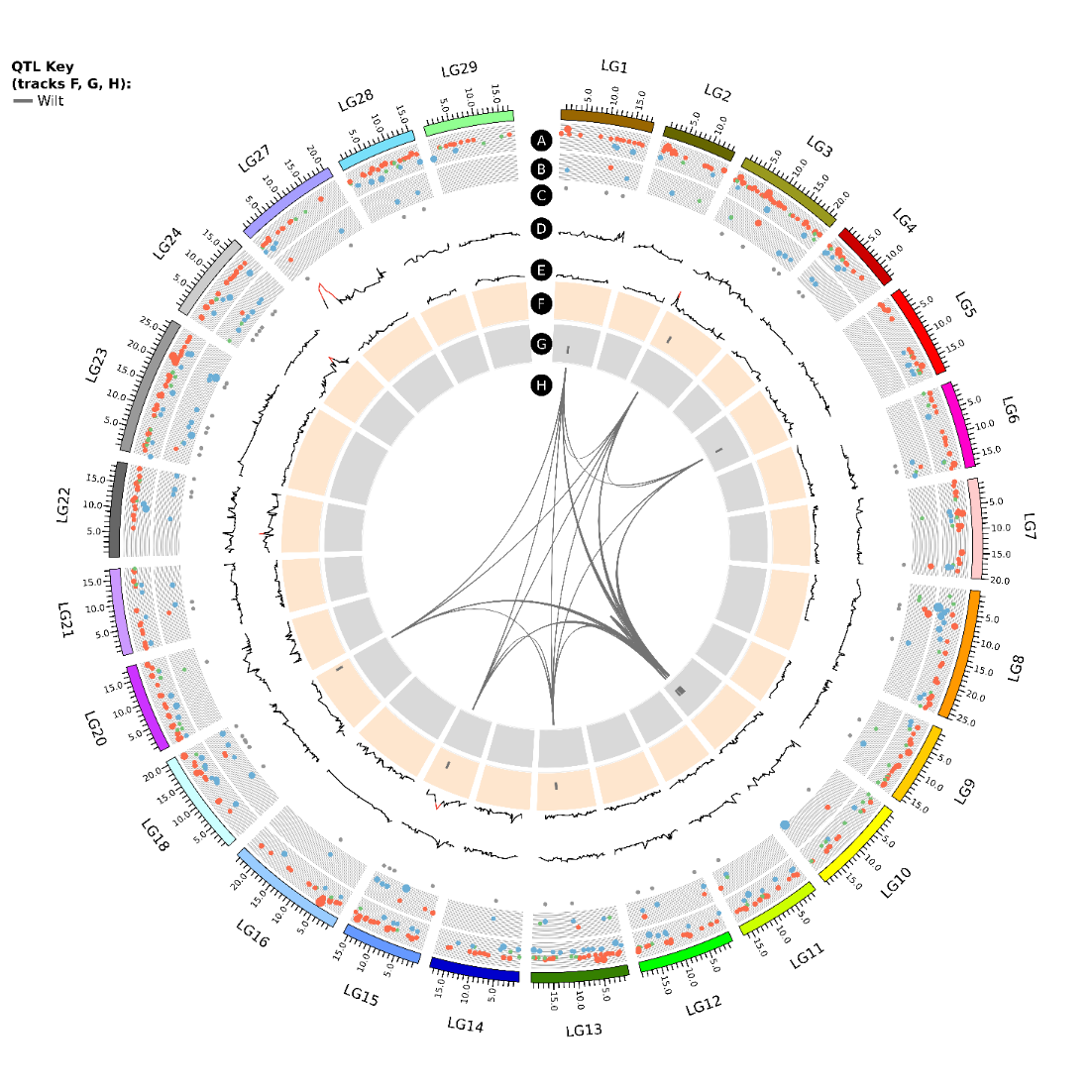

**Supplementary Figure 15a**. Circos plot of quantitative trait loci (QTLs) for Leaf_spots and Wilt phenotypes and RNA-seq data associated with *Psa* tolerant and susceptible genotypes, anchored on the chromosomes of the Red5 genome version 1.6.9 associated with QTLs. Tracks A and B represent differentially expressed gene (DEGs) with logFC ±2 and above in fully tolerant (Psa-FT) vs susceptible (Psa-Sus) and tolerant to medium tolerant (PsaTMT) vs Psa-Sus genotypes, respectively. A) blue circles are upregulated and red circles are downregulated genes in Psa-FT compared to Psa-Sus genotypes. B) blue circles are downregulated and red circles are upregulated genes in Psa-TMT compared to Psa-Sus genotypes. Genes with logFC between -1 and 1 are represented by green circles. Increase in circle diameter indicates increasing logFC value. C) DEGs common to Psa-FT and Psa-TMT. Track D and E are LOD values for Psa_score_Field for ‘Hort16A’ and P1 respectively. The lines change from black to red for a LOD score >3.Track F and G are QTLs detected from the phenotypes listed in QTL key  in ‘Hort16A’ and P1 respectively. H represents the lines connecting QTLs for similar phenotype on different chromosomes.

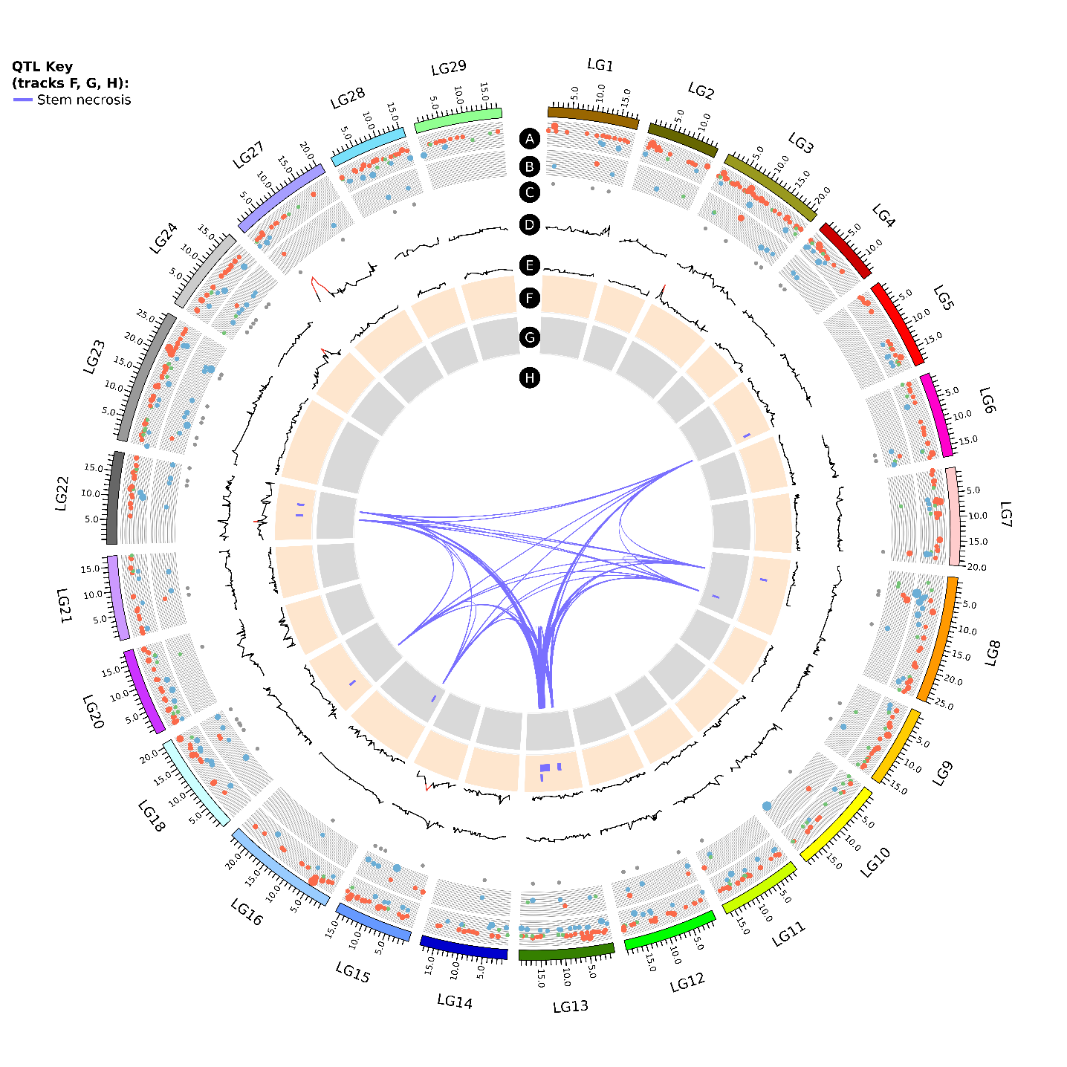

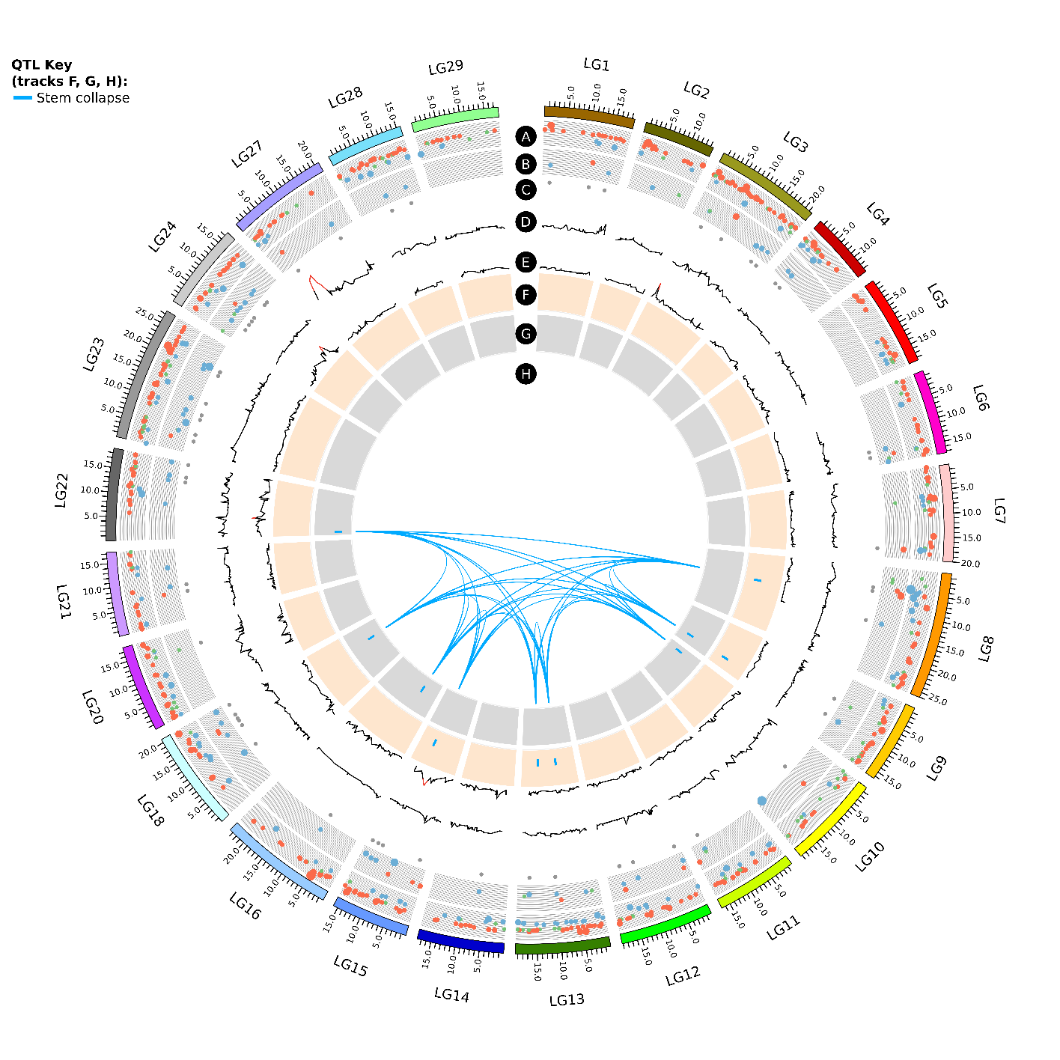

**Supplementary Figure 15b**. Circos plot of quantitative trait loci (QTLs) for Stem_necrosis and Stem_collapse phenotypes and RNA-seq data associated with *Psa* tolerant and susceptible genotypes, anchored on the chromosomes of Red5 genome associated with QTLs only. Track A and B represents DEGs with logFC ±2 and above in Psa-FT vs Psa-Sus and PsaTMT vs Psa-Sus, respectively. A) blue circles are upregulated and red circles are downregulated genes in FT compared to Sus genotypes. B) blue circles are downregulated and red circles are upregulated genes in TMT compared to Sus genotypes. Genes with logFC between -1 and 1 are presented by green circles. Increase in circle diameter indicates increasing logFC value. C) DEGs common to FT and TMT. Track D and E are LOD values for Psa_score_Field for ‘Hort16A’ and P1 respectively. The lines change from black to red above a LOD score of 3.Track F and G are QTLs detected from the phenotypes listed in QTL key in ‘Hort16A’ and P1 respectively. H represents the lines connecting QTLs for similar phenotype on different chromosomes.

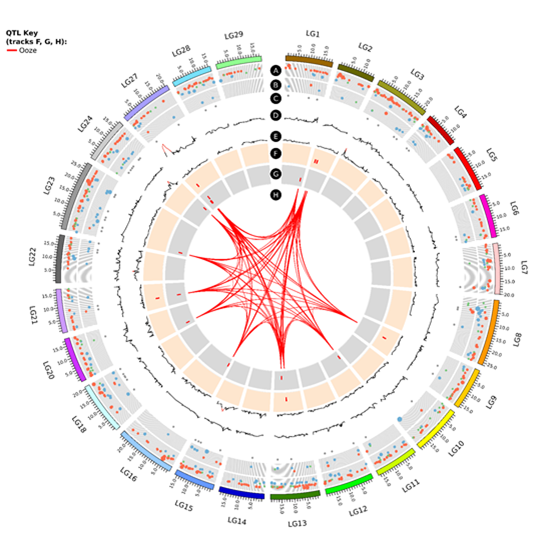

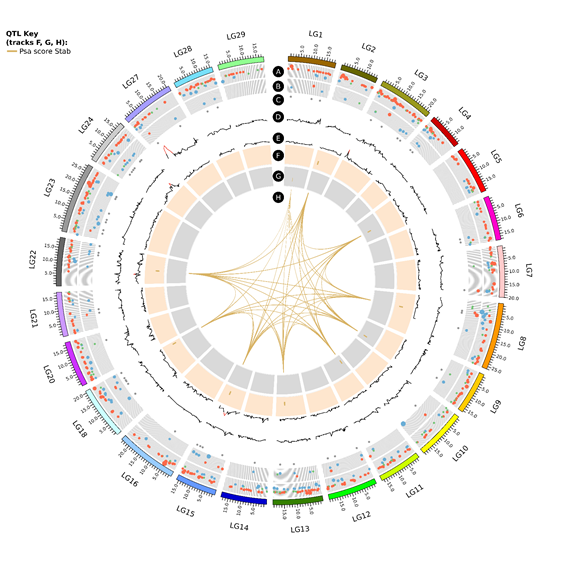

**Supplementary Figure 15c**. Circos plot of quantitative trait loci (QTLs) for Ooze and Psa_score_Stab phenotypes and RNA-seq data associated with *Psa* tolerant and susceptible genotypes, anchored on the chromosomes of Red5 genome associated with QTLs only. Track A and B represents DEGs with logFC ±2 and above in Psa-FT vs Psa-Sus and PsaTMT vs Psa-Sus, respectively. A) blue circles are upregulated and red circles are downregulated genes in FT compared to Sus genotypes. B) blue circles are downregulated and red circles are upregulated genes in TMT compared to Sus genotypes. Genes with logFC between -1 and 1 are presented by green circles. Increase in circle diameter indicates increasing logFC value. C) DEGs common to FT and TMT. Track D and E are LOD values for Psa_score_Field for ‘Hort16A’ and P1 respectively. The lines change from black to red above a LOD score of 3.Track F and G are QTLs detected from the phenotypes listed in QTL key in ‘Hort16A’ and P1 respectively. H represents the lines connecting QTLs for similar phenotype on different chromosomes.

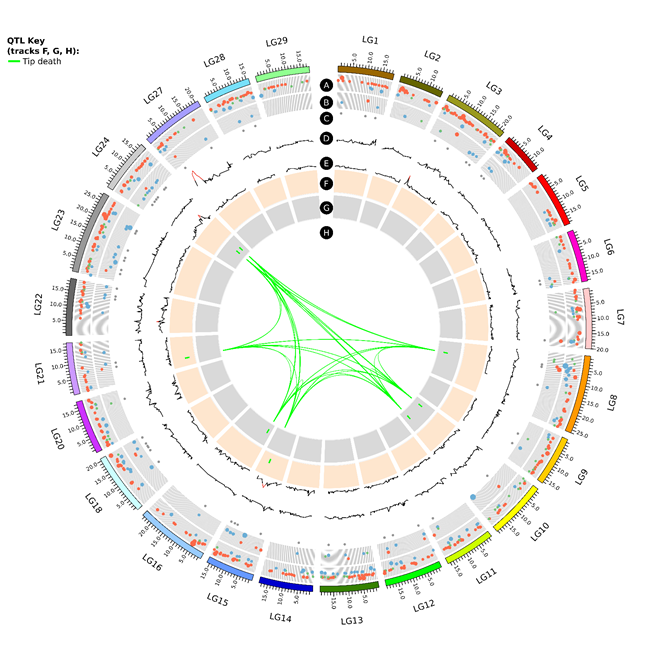

**Supplementary Figure 15d**. Circos plot of quantitative trait loci (QTLs) for Tip_death phenotype and RNA-seq data associated with *Psa* tolerant and susceptible genotypes, anchored on the chromosomes of Red5 genome associated with QTLs only. Track A and B represents DEGs with logFC ±2 and above in Psa-FT vs Psa-Sus and PsaTMT vs Psa-Sus, respectively. A) blue circles are upregulated and red circles are downregulated genes in FT compared to Sus genotypes. B) blue circles are downregulated and red circles are upregulated genes in TMT compared to Sus genotypes. Genes with logFC between -1 and 1 are presented by green circles. Increase in circle diameter indicates increasing logFC value. C) DEGs common to FT and TMT. Track D and E are LOD values for Psa_score_Field for ‘Hort16A’ and P1 respectively. The lines change from black to red above a LOD score of 3.Track F and G are QTLs detected from the phenotypes listed in QTL key in ‘Hort16A’ and P1 respectively. H represents the lines connecting QTLs for similar phenotype on different chromosomes.

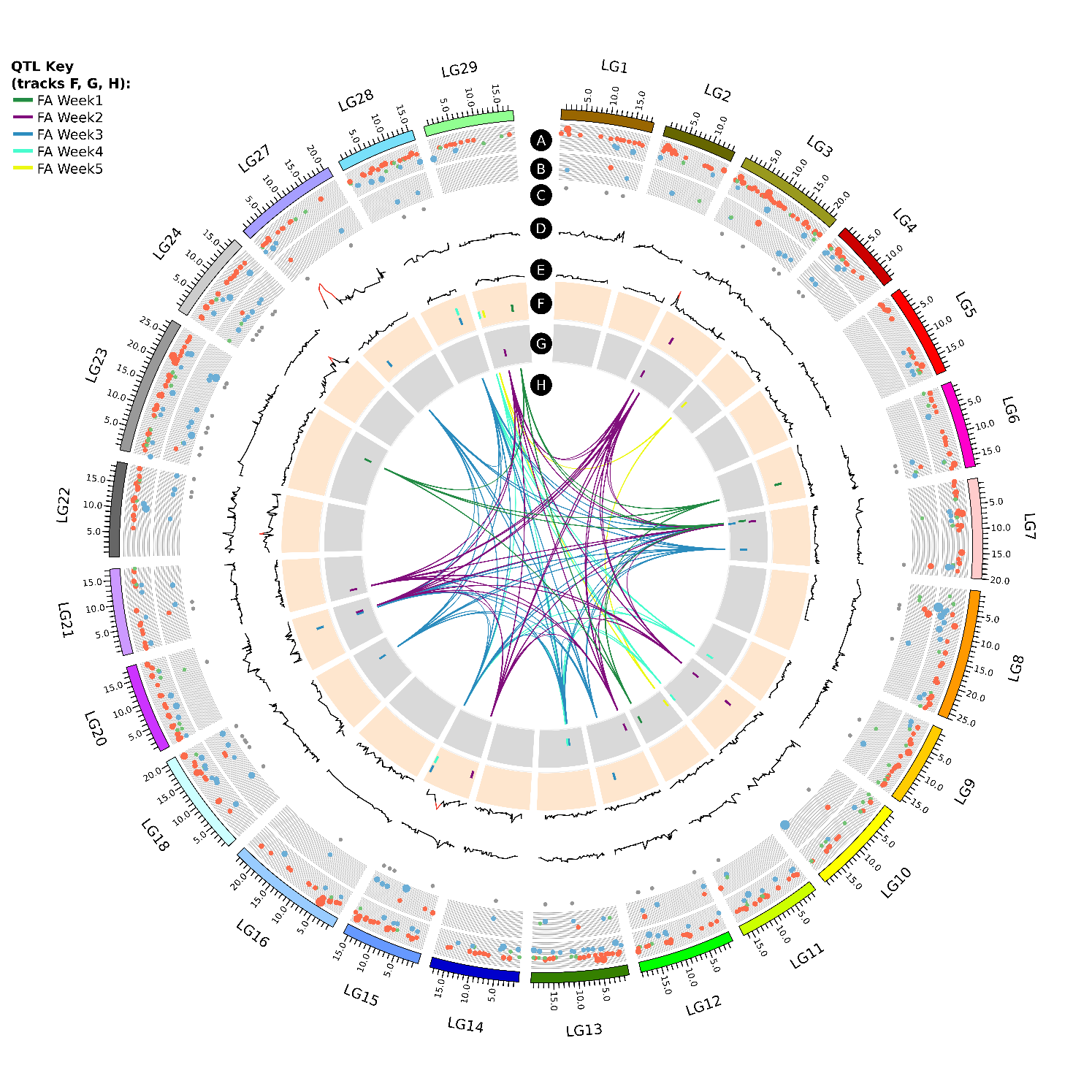

**Supplementary Figure 16**. Circos plot of QTLs for Flood assays (FA) phenotypes at time points of Week1-Week5 and RNA-seq data associated with *Psa* tolerant and susceptible genotypes, anchored on the chromosomes of Red5 genome associated with QTLs only. Track A and B represents DEGs with logFC ±2 and above in Psa-FT vs Psa-Sus and PsaTMT vs Psa-Sus, respectively. A) blue circles are upregulated and red circles are downregulated genes in FT compared to Sus genotypes. B) blue circles are downregulated and red circles are upregulated genes in TMT compared to Sus genotypes. Genes with logFC between -1 and 1 are presented by green circles. Increase in circle diameter indicates increasing logFC value. C) DEGs common to FT and TMT. Track D and E are LOD values for Psa_score_Field for ‘Hort16A’ and P1 respectively. The lines change from black to red above a LOD score of 3.Track F and G are QTLs detected from all phenotypes listed in QTL key in ‘Hort16A’ and P1 respectively. H represents the lines connecting QTLs for similar phenotype on different chromosomes.

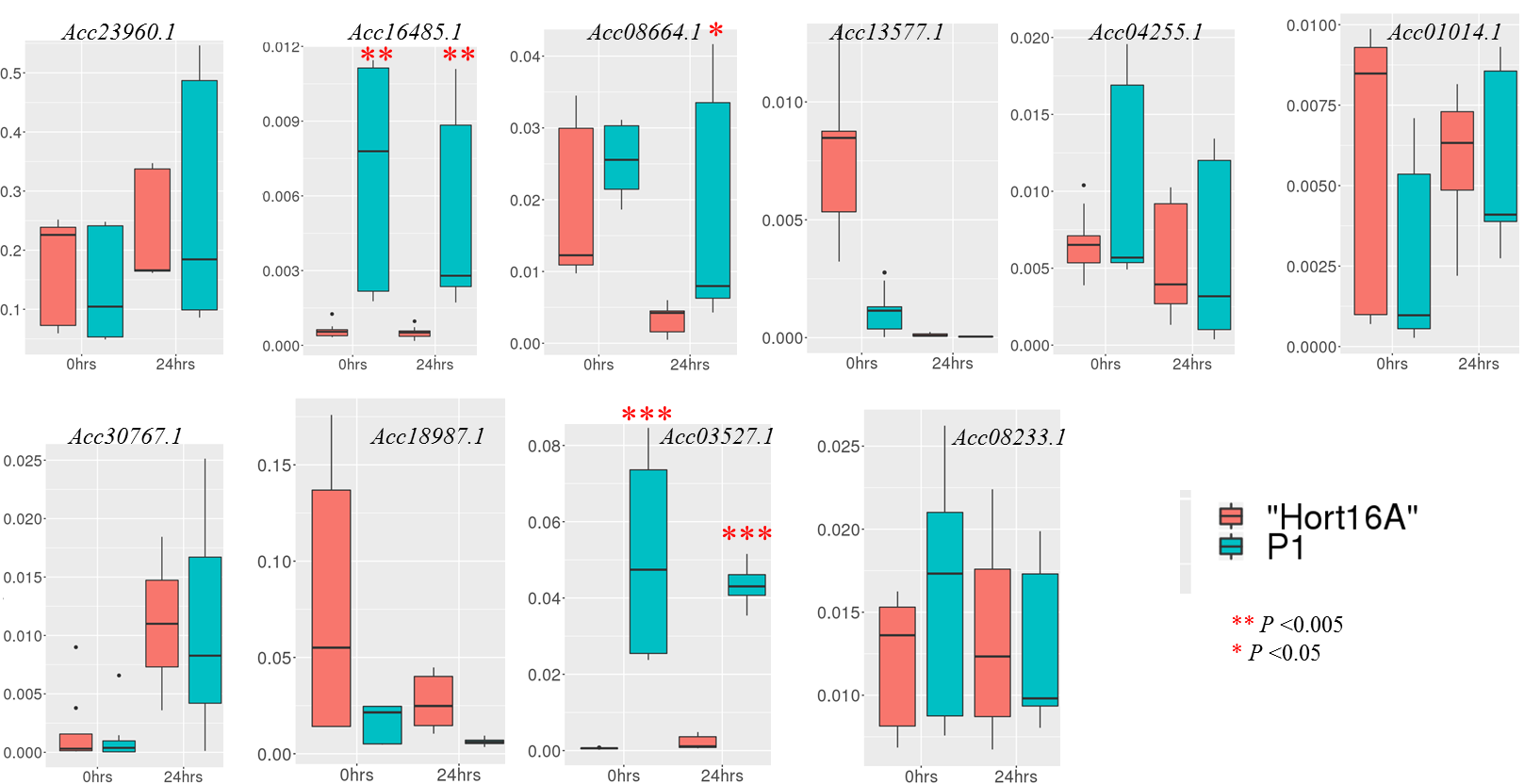

**Supplementary Figure 17**. Real-time quantitative RT-PCR of candidate genes in “Hort16A” and P1 post-Psa infection. Data represents gene expression of the candidate genes in three clonal replicates for each genotype at 0 and 24 hr post-infection and expression relative to the mean of the expression of *Actin* and *Ubiquitin* genes is plotted using geom_boxplot. Asterisks represent statistically significant differences in the relative expression of the candidate genes in P1 compared to “Hort16A” using Student’s *t*-test.

**Supplementary Table 1**. A) Kruskal-Wallis test for quantitative trait loci in ‘Hort16A’ and P1 based on phenotypic scores from the pilot trial. B) Phenotypic variance in Psa_score_Field explained by single nucleotide and multi-allelic genetic markers. Table represents outcome of linear regression analysis for the Psa_score_Field in August 2018 and genetic markers under each QTL as well as the combination of markers from QTLs. R^2^ (%) represents phenotypic variation, v.r. represents variance ratio for the F value and F pr. is the probability test for significance.

A)

| **‘Hort16A’** |  |  |  |
| --- | --- | --- | --- |
| Phenotype | Linkage group | Marker | *K* value |
| Cane_death | LG6 | S6_12039943 | 10 |
| Psa_score_Field | LG27 | S27_5573914 | 5 |
|  | LG11 | S11_11366770 | 7 |
| **P1** |  |  |  |
| Phenotype | Linkage group | Marker | K value |
| Cane_death | LG1 | S1_2162228 | 7 |
|  | LG12 | S12_3128909 | 12 |
|  | LG24 | S24_12974421 | 12 |
|  | LG25.2 | S25_19074112 | 6 |
| Tip_death | LG11 | S11_1487348, | 9 |
|  | LG24 | S24_12974421 | 14 |
|  | LG25.2 | S25_16016646 | 9 |
|  | LG27.1 | S27_4396077 | 9 |
| Ooze | LG27.1 | S27_2387628 | 9 |
|  | LG20 | S20_15144910 | 6 |
| Psa_score_Field | LG27.2 | Hy_sc27_5173960 | 9 |
|  | LG23 | S23_10035744 | 7 |
|  | LG28 | S28_1476180 | 7 |

B)

| Markers | R^2^ (%) | v.r. | F pr. | Type of markers |
| --- | --- | --- | --- | --- |
| LG27_G9P1 | 16.6 | 45.51 | <.001 | SNP |
| SSRLG27_439F4R4 | 19.5 | 18.9 | <.001 | SSR |
| LG14_E6P3 | 9.6 | 8.88 | <.001 | SNP |
| SSRLG22_8032664 | 4.6 | 4.48 | 0.004 | SSR |
| 3 QTLs combined LG27+LG14+LG22 | 37.5 | 3.24 | <.001 |  |
| 4 QTLs combined LG27+LG14+LG22+LG28 | 40.6 | 2.03 | <.001 |  |

**Supplementary Table 2**. Candidate genes in LG27 QTL.

| **Gene Ontology and function** | ***Actinidia* gene ID** |
| --- | --- |
| Cysteine rich receptor-like protein kinases | Acc30852.1, Acc30853.1, Acc30854.1 |
| Leaf rust disease-resistance receptor-like protein kinase | Acc30861.1 |
| LRR-receptor-like serine/threonine- protein kinase and receptor-like kinases | Acc30775.1, Acc30786.1, Acc30787.1, Acc30788.1, Acc30802.1, Acc31334.1 |
| Detoxification Proteins | Acc31349.1 and Acc31346.1 |
| Sugar transporter | Acc30857.1 |
| Glutamine dumper 1-like | Acc30812 |
| UDP-glycosyltransferase | Acc30767.1 |
| Alpha-1,4-glucan-protein synthase | Acc30791.1 |
| Glycosyltransferase | Acc30810.1 |
| Myo-inositol phosphotransferase | Acc31333.1 |
| GRAS – transcription factor | Acc30792.1 |
| MYB48– transcription factor | Acc30793.1 |
| Auxin-responsive protein | Acc31335.1 |
| Auxin transport protein | Acc30798.1 |
| Histone chaperone ASF1B | Acc31348.1 |
| Heat repeat-containing protein | Acc30840.1 |
| Cytochrome P450 85A1 | Acc30790.1 |
| EXO70E1 | Acc30773.1 |

**Supplementary Table 3**. Kruskal-Wallis test for quantitative trait loci (QTLs) in ‘Hort16A’ based on phenotypic scores from the stab bioassay. QTLs that overlap for control of different phenotypes are highlighted in same colour. The significance test employed oneway ANOVA with *P*-value < 0.05 = **; < 0.01 = ***; < 0.005 = ****; < 0.0001 = *****. GBS = genotyping by sequencing.

| Phenotype | Linkage group (LG) | Position (cM) | GBS Marker | K value | *P* value |
| --- | --- | --- | --- | --- | --- |
| Stem_necrosis | LG13 | 81.801 | S13_13629983 | 21.341 | ******* |
|  | LG15 | 65.029 | S15_8860868 | 8.766 | **** |
|  | LG18 | 79.033 | S18_5833359 | 9.274 | **** |
|  | LG22 | 77.151 | S22_12788245 | 8.529 | **** |
|  | LG5 | 28.204 | S5_17098703 | 10.202 | *** |
|  | LG8 | 77.675 | S8_8206774 | 12.664 | ****** |
| Ooze | LG2 | 30.063 | S2_2629030 | 10.831 | ***** |
|  | LG2 | 44.782 | S2_4100930 | 10.276 | **** |
|  | LG9 | 47.864 | S9_7227947 | 8.122 | **** |
|  | LG13 | 81.985 | S13_10678481 | 8.392 | **** |
|  | LG15 | 123.332 | S15_16622910 | 7.991 | **** |
|  | LG21 | 42.256 | S21_7135277 | 10.272 | *** |
|  | LG21 | 59.678 | HY21_9528792 | 8.324 | **** |
|  | LG27* | 80.87 | S27_4358305 | 6.148 | ** |
| Leaf_spots | LG2 | 29.568 | S2_2629585 | 13.889 | ****** |
|  | LG5 | 107.61 | S5_10632553 | 11.959 | **** |
|  | LG13 | 82.173 | S13_10678573 | 7.099 | *** |
|  | LG18 | 19.143 | S18_19172999 | 7.239 | *** |
| Wilt | LG3 | 92.768 | S3_5415275 | 9.111 | **** |
|  | LG7 | 74.379 | S17_14039379 | 8.514 | **** |
|  | LG11 | 98.208 | HY11_15736082 | 7.641 | *** |
|  | LG13 | 81.511 | S13_12912903 | 10.931 | **** |
|  | LG15 | 72.042 | S15_10684867 | 11.042 | ***** |
|  | LG18 | 19.143 | S18_19172999 | 18.15 | ******* |
| Tip_death | LG15 | 72.042 | S15_10684867 | 9.726 | **** |
|  | LG17 | 16.694 | S28_3242235 | 8.882 | **** |
|  | LG21 | 55.981 | S21_4644216 | 9.736 | **** |
|  | LG26 | 106.789 | S26_19872390 | 21.486 | ******* |
| Stem_collapse | LG7 | 74.776 | S17_14039198 | 7.079 | *** |
|  | LG8 | 81.673 | S8_9776918 | 7.181 | *** |
|  | LG9 | 70.784 | S9_12647445 | 9.474 | *** |
|  | LG13 | 65.222 | S13_7128533 | 12.252 | ****** |
|  | LG13 | 81.801 | S13_13629964 | 11.013 | ***** |
|  | LG15 | 72.042 | S15_10684867 | 7.05 | *** |
| Psa_score_Stab | LG2 | 29.568 | S2_2629585 | 11.664 | ***** |
|  | LG7 | 74.379 | S17_14039379 | 8.037 | **** |
|  | LG8 | 80.933 | S8_9536148 | 7.012 | *** |
|  | LG9 | 70.784 | HY9_7656534 | 11.293 | **** |
|  | LG13 | 81.801 | S13_13629964 | 16.503 | ******* |
|  | LG15 | 65.029 | S15_8860868 | 8.394 | **** |
|  | LG15 | 72.042 | S15_10684867 | 8.017 | **** |
|  | LG18 | 19.143 | S18_19172999 | 11.319 | ***** |

**Supplementary Table 4**. Kruskal-Wallis test for quantitative trait loci (QTLs) in P1 based on phenotypic scores from the stab bioassay. QTLs that overlap for control of different phenotypes are highlighted in same colour. The significance test employed oneway ANOVA with *P*-value < 0.05 = **; < 0.01 = ***; < 0.005 = ****; < 0.0001 = *****. GBS = genotyping by sequencing.

| Phenotype | Linkage group (LG) | Position(cM) | GBS Marker | K value | *P* value |
| --- | --- | --- | --- | --- | --- |
| Stem_necrosis | LG16.1 | 23.6 | S16_2880477 | 14.344 | ****** |
|  | LG8.2 | 1.782 | S8_19793836 | 7.239 | *** |
| Ooze | LG1 | 100.816 | S1_16292862 | 9.265 | **** |
|  | LG11.1 | 18.223 | S11_622686 | 9.135 | **** |
|  | LG13 | 84.998 | S13_13569603 | 11.228 | **** |
|  | LG16.1 | 2.884 | S16_398834 | 6.921 | *** |
|  | LG20.1 | 25.425 | S20_2567187 | 8.931 | **** |
|  | LG21.2 | 48.898 | S21_7135486 | 10.008 | *** |
|  | LG23.1 | 56.405 | S23_2231041 | 9.831 | **** |
|  | LG24 | 89.873 | S24_16024893 | 10.658 | **** |
|  | LG27.1* | 27.521 | S27_4621046 | 13.604 | ****** |
| Leaf_spots | LG1 | 49.483 | S8_2653760 | 8.966 | **** |
|  | LG5 | 44.586 | S5_10632553 | 11.959 | **** |
|  | LG7 | 23.045 | S7_2585468 | 9.576 | *** |
|  | LG15 | 33.881 | S15_11896346 | 10.631 | **** |
| Wilt | LG1 | 46.388 | S1_6558110 | 8.088 | **** |
|  | LG5 | 44.586 | S5_10632553 | 10.013 | *** |
|  | LG10 | 69.892 | S10_14907026 | 11.651 | **** |
|  | LG10 | 71.813 | HY30_1618648 | 18.979 | ******* |
|  | LG10 | 85.154 | S10_16015121 | 15.796 | ****** |
| Tip_death | LG8.1 | 54.543 | S8_8378507 | 8.892 | **** |
|  | LG10 | 33.985 | S10_3720973 | 9.842 | **** |
|  | LG10 | 79.942 | S10_15724230 | 7.921 | **** |
|  | LG16.1 | 21.789 | S16_2518111 | 9.614 | **** |
|  | LG24 | 62.528 | S24_11270794 | 8.688 | **** |
|  | LG24 | 94.111 | S24_14459417 | 13.719 | ****** |
| Stem_collapse | LG9 | 69.705 | S9_12647445 | 9.474 | *** |
|  | LG10 | 25.382 | S10_2439579 | 9.596 | **** |
|  | LG16.1 | 64.815 | S16_7213349 | 8.553 | **** |
|  | LG18 | 68.985 | S18_13967484 | 8.398 | **** |
|  | LG22 | 9.402 | S22_1422226 | 6.988 | *** |
| Psa_score_Stab | LG1 | 53.82 | S25_566827 | 12.709 | ****** |
|  | LG5 | 44.586 | S5_10632553 | 9.531 | *** |
|  | LG9 | 69.705 | S9_12647445 | 11.293 | **** |
|  | LG11.1 | 35.719 | S11_2346142 | 7.089 | *** |
|  | LG16.1 | 49.862 | HY16_5627871 | 8.617 | **** |
|  | LG23.1 | 81.863 | HY23_11642290 | 13.119 | ****** |

**Supplementary Table 5**. Kruskal-Wallis test for quantitative trait loci (QTLs) in ‘Hort16A’ and P1 based on phenotypic scores from flood assay. QTLs that overlap for control of different phenotypes are highlighted in same colour. The significance test employed oneway ANOVA with *P*-value < 0.05 = **; < 0.01 = ***; < 0.005 = ****; < 0.0001 = *****. GBS = genotyping by sequencing.

| ‘Hort16A’ |  |  |  |  |  |
| --- | --- | --- | --- | --- | --- |
| Phenotype | Linkage group (LG) | Position(cM) | GBS Marker | K value | *P* value |
| FA_Week1 | LG22 | 42.471 | S27_5310452 | 9.857 | *** |
|  | LG29 | 59.373 | S29_12351539 | 10.302 | **** |
|  | LG6 | 61.39 | S6_10837780 | 8.613 | **** |
| FA_Week2 | LG3 | 94.114 | S3_6302253 | 9.25 | *** |
|  | LG10 | 63.059 | S10_5024040 | 13.281 | **** |
|  | LG15 | 7.097 | S15_811591 | 8.159 | **** |
| FA_Week3 | LG12 | 67.455 | S12_13273020 | 7.625 | *** |
|  | LG15 | 113.564 | S15_15234101 | 8.299 | **** |
|  | LG20 | 46.98 | S20_12249726 | 12.391 | **** |
|  | LG27* | 65.587 | S27_4853516 | 6.923 | *** |
|  | LG28 | 67.438 | S28_11467605 | 11.44 | ***** |
| FA_Week4 | LG15 | 112.524 | S15_15233794 | 10.197 | **** |
|  | LG28 | 67.438 | S28_11467605 | 6.719 | *** |
|  | LG29 | 4.444 | S29_237182 | 8.139 | **** |
| FA_Week5 | LG6 | 116.351 | HY6_348464 | 10.756 | **** |
|  | LG29 | 20.567 | S29_1885874 | 10.687 | **** |
| P1 |  |  |  |  |  |
| FA_Week1 | LG7 | 22.24 | S7_2585362 | 7.101 | *** |
|  | LG10 | 44.995 | HY10_12312324 | 9.284 | **** |
|  | LG11.2 | 10.827 | S11_14629177 | 10.92 | **** |
|  | LG19.2 | 17.311 | S19_17614706 | 6.678 | *** |
|  | LG22* | 56.415 | S2_8357780 | 9.475 | **** |
|  | LG23.2 | 69.922 | S23_19456898 | 8.642 | **** |
| FA_Week2 | LG3.2 | 30.916 | S3_3107847 | 8.545 | **** |
|  | LG7 | 26.191 | S7_3201679 | 10.537 | **** |
|  | LG10 | 50.4 | S10_5024040 | 13.281 | **** |
|  | LG12 | 26.391 | S12_2655842 | 8.231 | **** |
|  | LG20.2 | 61.903 | S20_13104420 | 8.651 | **** |
|  | LG21.2 | 15.654 | S21_2673357 | 7.929 | **** |
|  | LG29 | 31.945 | S29_6544286 | 9.726 | **** |
| FA_Week3 | LG7 | 26.512 | S7_3201779 | 7.16 | *** |
|  | LG7 | 99.961 | S7_15056423 | 10.477 | *** |
|  | LG13 | 62.725 | S13_6476298 | 10.329 | **** |
|  | LG18 | 68.985 | S18_13967511 | 7.481 | *** |
|  | LG20.2 | 53.444 | S20_12249726 | 12.391 | **** |
| FA_Week4 | LG9 | 79.041 | S9_13780881 | 7.913 | **** |
|  | LG10 | 108.883 | S10_19063003 | 9.69 | *** |
|  | LG13 | 48.882 | S13_7200707 | 9.096 | **** |
| FA_Week5 | LG4 | 10.86 | S4_1083119 | 10.655 | **** |
|  | LG6 | 16.444 | HY6_1332956 | 12.371 | ****** |
|  | LG8.1 | 13.86 | S30_11496759 | 8.486 | **** |
|  | LG11.1 | 16.477 | S11_623069 | 8.229 | **** |

**Supplementary Table 6**. Primers used for the study.

A) Genetic mapping

| LG | Primer name | Primer sequence | SSR repeat/SNP in gene |
| --- | --- | --- | --- |
| **27** | LG27_4396125M13For3 | TGTAAAACGACGGCCAGTGCCATGCTTACCGTCAAGTG | (CA)4 |
| **27** | LG27_4396125Rev3 | GCTCAAAATCAACCTGATCG |  |
| **27** | LG27_4396125M13For4 | TGTAAAACGACGGCCAGTAAACTTCCCCAGCCATGAAG | (CT)20 |
| **27** | LG27_4396125Rev4 | TTTTTCGGGTTCTAAGACCA |  |
| **27** | LG27_4396125M13For5 | TGTAAAACGACGGCCAGTCGTACTGCGTTCCCGTACC | (GA)11 |
| **27** | LG27_4396125Rev5 | TCAAGGTATCGGCATCCATC |  |
| **27** | S27_3447729M13 For3 | TGTAAAACGACGGCCAGTGGAAGAAAATGTTGAGCAAGG | GA(14) |
| **27** | S27_3447729 Rev3 | CACACGCATCTTGTTTGTCC |  |
| **28** | S28_1523778M13 For5 | TGTAAAACGACGGCCAGTCATCTGATTTGTTTCGTTACAGAAG | (TA)13 |
| **28** | S28_1523778 Rev5 | TCTGGTGGATAATATACGCTTTACC |  |
| **22** | LG22_8032664M13For3 | TGTAAAACGACGGCCAGTCCGTTACGGTTCCACTTTTT | GA(17) |
| **22** | LG22_8032664Rev3 | TTCATCTGGAGAACGGGAAT | LG22 Tol associated with P1 |
| **27** | G9P1 For | GAGATGTATGTGACAAGGC | Acc30822.1 |
| **27** | G9P1 Rev | CGCTTCTGCTTCTTCATCTGTC | Acc30822.1 |
| **14** | E6P3 For | GGTGGAGTCATCGCCACGAGCATCG | Acc15766.1 |
| **14** | E6P3 Rev | AAGTCATTCCGCCGGCAAT | Acc15766.1 |

B) RT-qRT-PCR

| Primer name | Sequence |
| --- | --- |
| Actin F | GTTCCTGCCATGTATGTTGC |
| Actin R | CACACCATCACCAGAATCCA |
| Ubiquitin F | AGATCCAAGACAAGGAGGGA |
| Ubiquitin R | TGTTATAATCGGCCAGGGTG |
| *Acc23960.1 F* | TCCGAGTCCGTGTCTGTACT |
| *Acc23960.1 R* | TCGGGGCTGAAGGAAATTCC |
| *Acc16485.1 F* | CAACATATTAGCACGGCAGGC |
| *Acc16485.1 R* | TGCTGTTGCACCAAAGAGGA |
| *Acc30767.1 F* | ATCGATAAGGAAATCATAGAGCATGATG |
| *Acc30767.1 R* | GAGCTGAGTTGCAAATCAGCCAGTC |
| *Acc08664.1 F* | GCACCCTACGCGGCTAATAT |
| *Acc08664.1 R* | CTGCCCCTGGAGTAATGCAA |
| *Acc18987.1 F* | TGCTGAACATGCCTGCAGAGAAAGCATGGGA |
| *Acc18987.1 R* | AGTCGGATACTTCCAGGGCTACCATCT |
| *Acc03527.1 F* | AGATGAAGAGAATCGAGAACCCCACGAGC |
| *Acc03527.1 R* | CAGCCACTTGGGCATCACAAAGAACGGA |
| *Acc08233.1 F* | TCATATCCATCATTGCTCTCCGATT |
| *Acc08233.1 R* | CTCCTCGATCCGTACAGTGAGATA |
| *Acc01014.1 F* | TCCCCGACAGAAGTGAAACG |
| *Acc01014.1 R* | TCTGCAACAGCTCTCATGCA |
| *Acc24057.1 F* | GTGACGTCATCTCCCTCGAC |
| *Acc24057.1 R* | TGCCCTGGGAGTGATCACTA |
| *Acc04255.1 F* | TTTCATGGAGTGGGCCACAG |
| *Acc04255.1 R* | CCCGGTGTCACTGAAATGGA |
| *Acc13577.1 F* | TGAGGAATACGCAGCCCAAC |
| *Acc13577.1 R* | TTTCCCGGAAAAGCCTGTCG |
