## Supplementary material for "Multiple quantitative trait loci contribute tolerance to bacterial canker incited by *Pseudomonas syringae* pv. *actinidiae* in kiwifruit (*Actinidia chinensis*)": Table 1

**Table 1. Candidates from differentially expressed genes in field tolerant genotypes.** Psa-tolerant plants (FT), Psa Tolerant to Medium Tolerant /Psa-TMT and Psa-susceptible genotypes (Psa-Sus).

|  | **Gene Ontology and function** | ***Actinidia* gene ID** | ***Arabidopsis* orthologue** |
| --- | --- | --- | --- |
| **Upregulated in TMT and FT** | MATE efflux family protein, Protein detoxification | Acc00747.1 | AT5G52450.1/*DTX16* |
|  | *MADS*-box | Acc03527.1 | AT5G62165.2/AGL42 |
|  | Terpene synthases | Acc13740.1, Acc13742.1, Acc22685.1, Acc22685.1 | AT5G23960.2 /TPS21 |
|  | Major Latex Protein (MLP)-like protein | Acc18987.1, Acc13742.1 | AT1G24020.1/MLP28 |
|  | Thioredoxin-like protein | Acc20584.1, Acc20586.1 | AT1G11530.1/CXXS1 |
|  | Cellulose synthase-like protein | Acc27502.1, Acc15562.1 | AT4G24010.1/CSLG1 |
|  | *UDP-glycosyltransferase* | Acc30767.1 | AT3G02100.1/UGT72B1 |
|  | WD40-repeat containing super-family protein | Acc23960.1 | AT1G78070.1 |
|  | Protein of unknown function, UV-B*-*induced protein, *DUF760* | Acc25706.1, Acc14728.1 | AT3G07310.1 |
|  | Protein of unknown function, *DUF247* | Acc08761.1 | AT4G31980.1 |
|  | Ammonium transporter | Acc08664.1 | AT2G38290.1/AMT2 |
|  | *Chloroplastic, 3-ketoacyl-acyl carrier protein synthase* | Acc08233.1 | AT1G24360.1/KASI |
|  | *Alpha-glucan phosphorylase* | Acc16485.1 | AT3G46970.1 |
| **Downregulated in TMT and FT** | *Histone superfamily protein, Histone H2A*, Chromatin assembly factor-1 | Acc15097.1, Acc15099.1, Acc17300.1, Acc16944.1, Acc17279.1, Acc20675.1, Acc20918.1, Acc21661.1, Acc25126.1, Acc25392.1, Acc25885.1, Acc26149.1, Acc26150.1, Acc26360.1, Acc27699.1, Acc30085.1, Acc30211.1, Acc30253.1, Acc31646.1, Acc32318.1 | AT1G09200.1, AT1G65470.1, AT1G65470.1, AT2G28720.1, AT4G27230.1, AT5G59910.1, AT5G02560.1, AT3G45930.1, AT5G22650.2, AT1G54690.1 |
|  | *Salicylate carboxymethyltransferase* | Acc01014.1 | AT1G19640.1 |
|  | Auxin efflux carrier family protein | Acc24057.1 | AT1G77110.1 |
|  | Acyl-CoA N-acyltransferases (NAT) superfamily protein | Acc04255.1 | AT2G32030.1 |
|  | Serine/threonine-protein kinase PBS1-like | Acc17448.1 | AT3G20530.1 |
|  | Pathogenesis-related thaumatin superfamily protein | Acc25881.1 | AT2G28790.1 |
